# Free Fatty Acid Receptor 2 Allosterism is Defined By Cellular Context

**DOI:** 10.64898/2025.12.18.695082

**Authors:** Simon Lind, Ayaan Abdi Ali, Sarah Al Hamoud Al Asswad, Volker M. Lauschke, Huamei Forsman, Claes Dahlgren, Linda Johansson

## Abstract

**Background:** Allosteric modulators offer a way to fine-tune GPCR signaling in the presence of endogenous ligands. The short-chain fatty acid receptor FFA2R (GPR43) recognizes propionate and allosteric ligands such as Cmp58 and AZ1729. We characterized FFA2R signaling and allosteric modulation using multiple cell models including HEK293, HL60 cells and primary human neutrophils.

**Methods:** FFA2R activation was assessed using complementary assays in HEK293 and HL60 cells as well as primary human neutrophils. G protein activation and β-arrestin recruitment were profiled using ebBRET biosensors. Ca^2+^ mobilization was measured with Fura-2, and reactive oxygen species (ROS) generation was quantified by isoluminol chemiluminescence. Pharmacological tools included the FFA2R antagonist CATPB, the Gα_q_ inhibitor YM-254890, and pertussis toxin (PTX).

**Results:** Propionate activated all tested G proteins except Gα_12_ in HEK293 cells and recruited both β-arrestin1 and β-arrestin2. The allosteric ligands Cmp58 and AZ1729 behaved as pathway-selective ago-PAMs. Alone they engaged a limited subset of G proteins with minimal β-arrestin recruitment, whereas in the presence of propionate they selectively potentiated Gα_i1_ while attenuating Gα_q/11_ and Gαi_2/3_. Fura-2 measurements coupled to YM-254890 treatment established that FFA2R couples to Gα_q_-dependent Ca^2+^ mobilization in HEK293 cells; Cmp58, but not AZ1729, enhanced Ca^2+^ responses at submaximal propionate concentrations. In primary neutrophils, propionate elicited Ca^2+^ transients but did not trigger NADPH oxidase–dependent ROS on its own. Either Cmp58 or AZ1729 enabled propionate-driven ROS, and their combination produced robust ROS. Transient FFA2R expression in HL60 cells reconstituted this neutrophil-like functional profile, including allosteric activation of ROS.

**Conclusions:** Across systems, Ca^2+^ mobilization emerged as a conserved, receptor-proximal output of FFA2R, while allosteric modulation by Cmp58 and AZ1729 promoted Gα_i/o_-biased signaling that enabled ROS generation. These data define pathway-selective allosterism at FFA2R and highlight Ca^2+^ mobilization and ROS as informative readouts for therapeutic strategies that exploit allosteric control.

## Introduction

G protein-coupled receptors (GPCRs) constitute the largest class of cell-surface signaling proteins^1^. Activated by diverse stimuli such as odors, fatty acids, and hormones, they trigger intracellular cascades that regulate key physiological processes. Across cell types, GPCRs are central to many diseases and thus serve as major drug targets, with approximately one-third of all approved drugs acting *via* GPCRs^2,3^. Understanding how ligands and heterotrimeric G proteins shape receptor signaling is critical for therapeutic design.

A key concept in GPCR pharmacology is allosteric modulation, in which ligands bind to receptor sites distinct from the orthosteric pocket to alter receptor activity^4,5^. Allosteric ligands can enhance (positive allosteric modulators, PAMs) or inhibit (negative allosteric modulators, NAMs) a receptor’s response to an orthosteric agonist^6,7^. Some allosteric ligands also display intrinsic agonism, and others combine agonism with modulation of an orthosteric agonist’s potency or efficacy (ago-allosteric modulators). Structural and functional studies reveal that many GPCRs can accommodate multiple allosteric sites, allowing distinct signaling outcomes not accessible through orthosteric ligands^8–11^. Because allosteric modulators can fine-tune receptor responses with high subtype selectivity and without competing with endogenous ligands, they are considered promising candidates for next-generation therapeutics^12–14^. Allosteric effects can appear markedly different depending on the cellular system examined, since each model highlights a distinct layer of receptor signaling. Systematic side-by-side comparisons, as performed here for FFA2R, are therefore essential to reveal conserved features of allosterism as well as context specific equivalents.

The free fatty acid receptor 2 (FFA2R; GPR43) is a short-chain fatty acid (SCFA) receptor activated by metabolites such as acetate, propionate, and butyrate derived from gut microbial fermentation^15,16^. Beyond serving as an energy source for intestinal cells, SCFAs act as endogenous ligands for FFA2R, mediating a range of physiological effects^7,8,13,17^. Through FFA2R, SCFAs shape innate immune responses, reinforce epithelial barrier integrity, modulate gut hormone secretion, and influence systemic metabolic and inflammatory set points.

In addition to these endogenous FFA2R orthosteric agonists, several allosteric ligands such as Cmp58^18,19^ and AZ1729^7,11,20^ have also been identified and characterized in different experimental systems. FFA2R is expressed in enteroendocrine cells, adipose tissue, and immune cells, including neutrophils^16,20,21^.

Although neutrophils are easy to isolate from peripheral human blood in large numbers, these cells are non-dividing with low protein synthesis and a short lifespan, which severely limits genetic manipulation and complicates detailed GPCR signaling studies^22,23^. To address this, we used HEK293 and HL60 cells as complementary models to dissect FFA2R signaling. The HEK293 cell line is a well-established model for GPCR research because of its efficient transfection, robust receptor expression, and suitability for real-time biosensor assays^24^. The enhanced bystander BRET (ebBRET) system measures G protein activation and β-arrestin recruitment in live cells, enabling quantification of pathway-selective signaling^25,26^. Because HEK293 cells lack endogenous FFA2R, transient transfection allows the analysis of receptor-specific responses to orthosteric and allosteric ligands.

The human leukemia cell line, HL60, has long been used as a model for functional and signaling studies that are difficult to perform in freshly isolated neutrophils^27^. These cells can be differentiated into neutrophil-like cells that reproduce some of the key neutrophil GPCR regulated effector functions including a functional reactive oxygen species (ROS) generating NADPH oxidase^28,29^. Like HEK293, HL60 cells lack endogenous FFA2R expression, allowing controlled receptor expression and functional assessment of an FFA2R-driven rise in the cytosolic concentration of free calcium ion (Ca^2+^) and the production ROS.

Herein, we investigated how cellular context shapes FFA2R allosterism and pathway selectivity. We compared responses to the endogenous ligand propionate and the xenobiotic allosteric ligands Cmp58 and AZ1729 across HEK293, HL60, and primary human neutrophils. We combined ebBRET profiling of G-protein activation and β-arrestin recruitment in HEK293 with functional readouts directly relevant to immune physiology. Ca^2+^ mobilization, measured with Fura-2, provided an early receptor-proximal signal in all three systems. ROS generation, quantified by isoluminol chemiluminescence, served as a myeloid-specific effector in neutrophils and FFA2R-transfected differentiated HL60 cells.

In HEK293 cells, propionate activated several G-protein pathways and recruited both β-arrestin 1 and 2, whereas Cmp58 and AZ1729 showed more restricted signaling and minimal β-arrestin effects. Across systems, FFA2R consistently coupled to Gα_q_-dependent Ca^2+^ mobilization, while robust ROS formation in neutrophils and differentiated HL60 cells required allosteric modulation. The combination of Cmp58 and AZ1729 produced cooperative ROS activation only in myeloid cells, pointing to a context-dependent, Gα_i/o_-linked mechanism.

Together, this comparative design establishes Ca^2+^ mobilization as a conserved output of FFA2R signaling across model systems and positions ROS formation as a context-dependent endpoint that reveals how orthosteric and allosteric ligands jointly shape receptor function.

## Results

### The propionate-activated free fatty acid receptor 2 (FFA2R) recruits most G protein subtypes

GPCRs can engage multiple signaling pathways or, conversely, display pathway selectivity, a phenomenon termed biased signaling^27,30^. The use of BRET-based effector membrane translocation assays provides an effective approach for profiling GPCR signaling by monitoring Gα protein activation and β-arrestin recruitment in living cells (**Fig. 1A**)^25,26^. Since terminally differentiated neutrophils exhibit restricted protein synthesis^22^, they are unsuitable for this assay. Therefore, HEK293 cells, which allow efficient transfection and robust receptor expression, were first used to study FFA2R signaling^24^. This system has also been successfully applied to investigate the related acid-sensing GPCR, HCAR1^31^.

**Fig.1:**
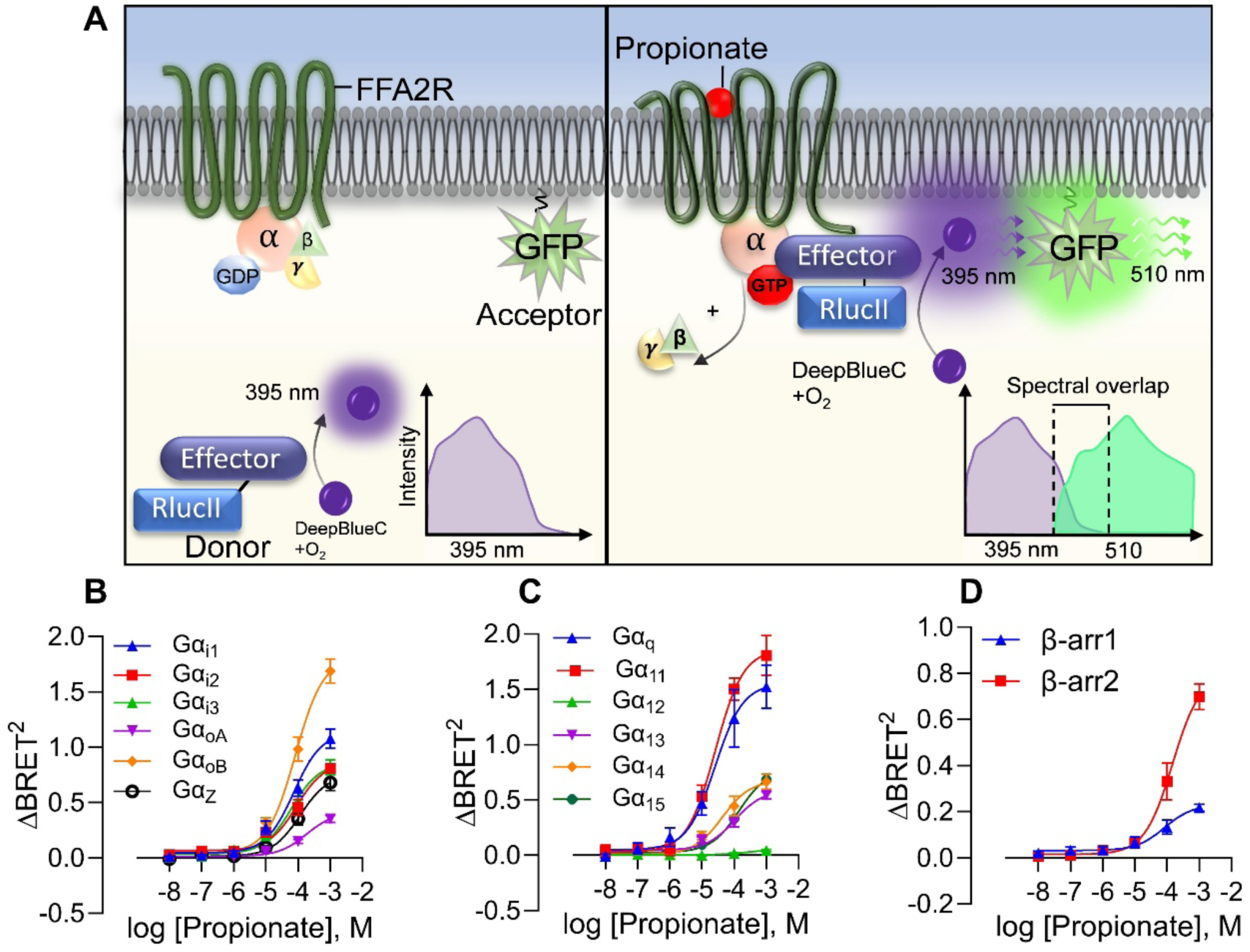
ebBRET profiling reveals endogenous ligand propionate to be a promiscuous G protein activator for FFA2R. **(A)** Overview of the effector membrane translocation assay (EMTA) used in this study to assess FFA2R-mediated G protein activation. Left panel: In the absence of ligand, the donor (*e.g.,* effector protein Rap1Gap to look at Gα_i/o_ activation) is not in close proximity and cannot excite the acceptor (membrane anchored GFP) resulting in emission of violet light (395 nm). Right panel: Agonist binding to FFA2R triggers G protein activation and dissociation of the G protein and subsequently the donor binding to the activated G protein. The close proximity between donor and acceptor promotes rGFP excitation and both violet and green light (510 nm) are emitted and the degree of spectral overlap is monitored in ebBRET. (**B**) Concentration-response curves of FFA2R endogenous agonist propionate monitoring the activation profile of the Gα_i/o_ subfamily. (**C**) Concentration-response curves FFA2R endogenous agonist propionate monitoring the activation profile of the Gα_q/11_, Gα_12/13_ and the Gα_14/15_ subfamily. This is performed using the effector membrane translocation assay (EMTA) with different biosensors monitoring the movement to the plasma membrane following ligand addition using rGFP-CAAX. (**D**) Concentration-response curves of FFA2R endogenous agonist propionate monitoring the β-arrestin recruitment for both β-arrestin1 and 2. (**B-D**) subfigures present nonlinear fit dose concentration of the compound in log (molar) on the x-axis and the unitless ΔBRET^2^ signal on the y-axis of at least three biological replicates performed in duplicates (n=3).

The orthosteric FFA2R agonist propionate induced a concentration-dependent activation of FFA2R expressed in HEK293T cells, an activation resulting in recruitment of Gα proteins belonging to different subtypes (**Fig. 1B**, **Table 1**). Propionate required relatively high concentrations to activate FFA2R in HEK293T cells and reached its maximal measurable response at 1 mM, the highest concentration that could be added without affecting cell viability (Supplementary Fig. S1A). In our assay, propionate produced clear activation across all Gα_i/o_ subtypes studied (**Fig. 1B**). Beyond the Gα_i/o_ family, propionate-activated FFA2R engaged every tested G protein pathway except Gα_12_ (**Fig. 1C**). Propionate also induced recruitment of β-arrestin1 and β-arrestin2 (**Fig. 1D**).

**Table 1:** EC_50_ values and max responses observed at concentration tested (ΔBRET^2^) for propionate using ebBRET. “–“ indicates no activity in that pathway.

| ebBRET<br>measurement | Propionate |  |
| --- | --- | --- |
|  | EC<sub>50</sub> (μM) | Max response |
| Gα<sub>i1</sub> | 21.4 | 0.7552 |
| Gα<sub>i2</sub> | 88.98 | 0.8652 |
| Gα<sub>i3</sub> | 61.89 | 0.8587 |
| Gα<sub>q</sub> | 25.56 | 1.554 |
| Gα<sub>11</sub> | 25.9 | 1.82 |
| Gα<sub>oA</sub> | 182.8 | 0.4152 |
| Gα<sub>oB</sub> | 81.8 | 1.812 |
| Gα<sub>z</sub> | 115.2 | 0.7568 |
| Gα<sub>12</sub> | - | - |
| Gα<sub>13</sub> | 103.7 | 0.5877 |
| Gα<sub>14</sub> | 42.58 | 0.6768 |
| Gα<sub>15</sub> | 140.3 | 0.68 |
| βarr1 | 83.59 | 0.23 |
| βarr2 | 147.5 | 0.7987 |

### The FFA2R ligands Cmp58 and AZ1729 alone activate the receptor to recruit fewer Gα protein subtypes and show decreased β-arrestin recruitment

The FFA2R specific ligands Cmp58 and AZ1729 have previously been characterized as ago-allosteric agonists (ago-PAMs) and as true positive allosteric modulators (PAMs) depending on the cell type^8,17,20,21^. Using the ebBRET approach in FFA2R-expressing HEK293T cells, both ligands activated the receptor on their own but engaged a more limited set of G-protein pathways compared to propionate, and each produced only minimal β-arrestin recruitment (**Fig. 2A–C**). Cmp58 and AZ1729 also generated measurable signals on Gα_q_ and Gα_15_ activation within the same assay framework (**Fig. 2A–C**; Supplementary Fig. S1C–D; **Table 2**). To functionally verify Gα_i/o_ coupling downstream of FFA2R, we quantified ligand effects on cAMP using the GloSensor cAMP assay. In HEK293T cells transiently expressing FFA2R, propionate, Cmp58, and AZ1729 independently produced concentration-dependent inhibition of forskolin-stimulated cAMP (Supplementary Fig. S1B).

**Fig.2:**
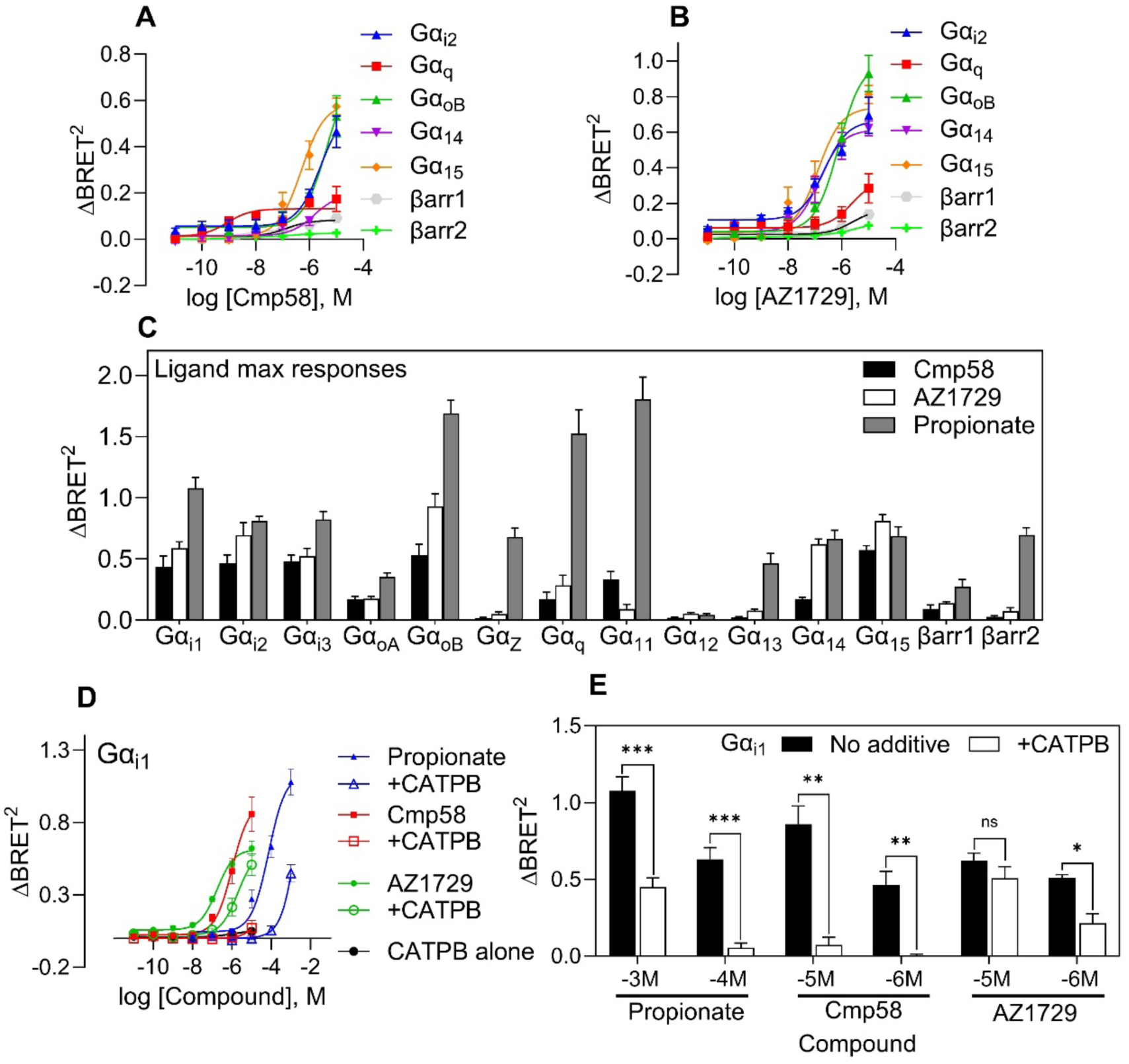
The two allosteric agonist Cmp58 and AZ1729 alone activate G proteins alone with minimal β-arrestin recruitment. (**A**) Concentration-response curves of FFA2R allosteric agonist Cmp58 monitoring the activation profile of the Gα_i2_, Gα_q_, Gα_oB,_ Gα_14/15_ subfamily as well as recruitment of β-arrestin1 and 2. (**B)** Same experimental setup as in **(A)** but the activating ligand is instead the allosteric agonist AZ1729. (**C**) Bar plots showing maximum ΔBRET2 responses (Maximum responses at the concentration tested) of Cmp58 (black bar), AZ1729 (white bar) or propionate (grey bar) for the different G-proteins as well as β-arrestin1 and 2 to visualize responses of ΔBRET^2^ of the different ebBRET biosensors used to study FFA2R signaling. The bar graph represents data from three independent experiments (mean+SEM); n = 3. **(D)** Dose dependent response curves of propionate (blue), Cmp58 (red), or AZ1729 (green) in the absence or presence of FFA2R antagonist CATPB (1 μM, 5 min pre incubation marked as hollow symbols in respective color) monitoring the activation profile of the Gα_i1_. The activation profile of CATPB (black line) alone was also followed. **(E)** Bar chart represents the (C) data, and the statistical analysis was performed using a paired Student’s *t* test comparing the peak ΔBRET^2^ responses for propionate, Cmp58, or AZ1729 at different concentration in the absence (black bars) and presence of the FFA2R antagonist CATPB (1 μM, white bars). The statistical analysis was performed using paired Student’s t-test comparing the peak responses in the absence and presence of CATPB. (mean ± SEM n = 3). (**A, B, E)** subfigures present nonlinear fit dose concentration of the compound in log (molar) on the x-axis and the unitless ΔBRET^2^ signal on the y-axis of at least three biological replicates performed in duplicates (n=3). All other results are from at least n=3 (technical duplicates).

**Table 2:** EC_50_ values and max responses observed at concentration tested (ΔBRET^2^) for Cmp58 and AZ1729 using ebBRET. “–“ indicate no activity in that pathway. N/A means data not available.

| ebBRET measurement | Cmp58 |  | AZ1729 |  | Cmp58+ AZ1729 |  |
| --- | --- | --- | --- | --- | --- | --- |
|  | EC<sub>50</sub> (nM) | Max response | EC<sub>50</sub> (nM) | Max response | EC<sub>50</sub> (nM) | Max response |
| Gα<sub>i1</sub> | 2282 | 0.5246 | 41 | 0.5839 | 70 | 0.6 |
| Gα<sub>i2</sub> | 2507 | 0.567 | 224 | 0.6645 | 235 | 0.5895 |
| Gα<sub>i3</sub> | 2106 | 0.5741 | 53 | 0.5534 | 50 | 0.4850 |
| Gα<sub>q</sub> | 0.87 | 0.1316 | 2391 | 0.3365 | 0.6413 | 0.1208 |
| Gα<sub>11</sub> | 10 039 | 0.33 | - | - | - | - |
| Gα<sub>oA</sub> | 4477 | 0.2373 | 186 | 0.1901 | N/A | N/A |
| Gα<sub>oB</sub> | 4378 | 0.7416 | 763 | 0.9936 | 916.6 | 0.9370 |
| Gα<sub>z</sub> | - | - | - | - | N/A | N/A |
| Gα<sub>12</sub> | - | - | - | - | - | - |
| Gα<sub>13</sub> | - | - | - | - | - | - |
| Gα<sub>14</sub> | 1494 | 0.1951 | 143 | 0.6144 | N/A | N/A |
| Gα<sub>15</sub> | 503 | 0.5877 | 131 | 0.7395 | 7465 | 0.4296 |
| βarr1 | 141 | 0.08271 | 2536 | 0.1662 | N/A | N/A |
| βarr2 | - | - | - | - | - | - |

Cmp58 and AZ1729 have been shown to bind to two different allosteric sites distinct from the orthosteric binding site^11,20^ but can still be inhibited by the orthosteric FFA2R antagonist CATPB^32^. Accordingly, CATPB inhibited the responses induced by propionate as well as by Cmp58, and AZ1729 (**Fig. 2D, E**). Inhibition was partial at the highest ligand concentrations but in accordance with competitive binding dynamics, the inhibitory effect of the antagonist was more pronounced when the concentrations of the activating ligands were reduced.

### The allosteric effect of Cmp58 and AZ1729 on the propionate-induced response is biased in HEK293 cells

The ebBRET experiments described above demonstrated that Cmp58 and AZ1729 each exhibit agonistic activity at FFA2R under these assay conditions. However, in both neutrophils and in HEK293T cells there is evidence of positive allosteric modulation^7,19,33^. The effects of a fixed concentration (1 µM) of the two ago-PAMS, respectively, on the propionate concentration-dependent recruitment of the different G proteins were determined. Both modulators enhanced the recruitment of Gα_i1_ to the propionate activated FFA2R (**Fig. 3A-B**). To note: vehicle signals and responses to Cmp58 or AZ1729 alone were subtracted, allowing the curves to reflect the net allosteric effect of propionate in the presence of each modulator (see Methods for details). Although, both Cmp58 and AZ1729 positively modulated the Gα_i1_ propionate response, they had low to slight negative effect on the recruitment of the other Gα_i/o_ subtypes Gα_i2_ and Gα_i3_ (**Fig. 3C–F**, **Table 3**, Supplementary Fig.S2A-F).

**Fig.3:**
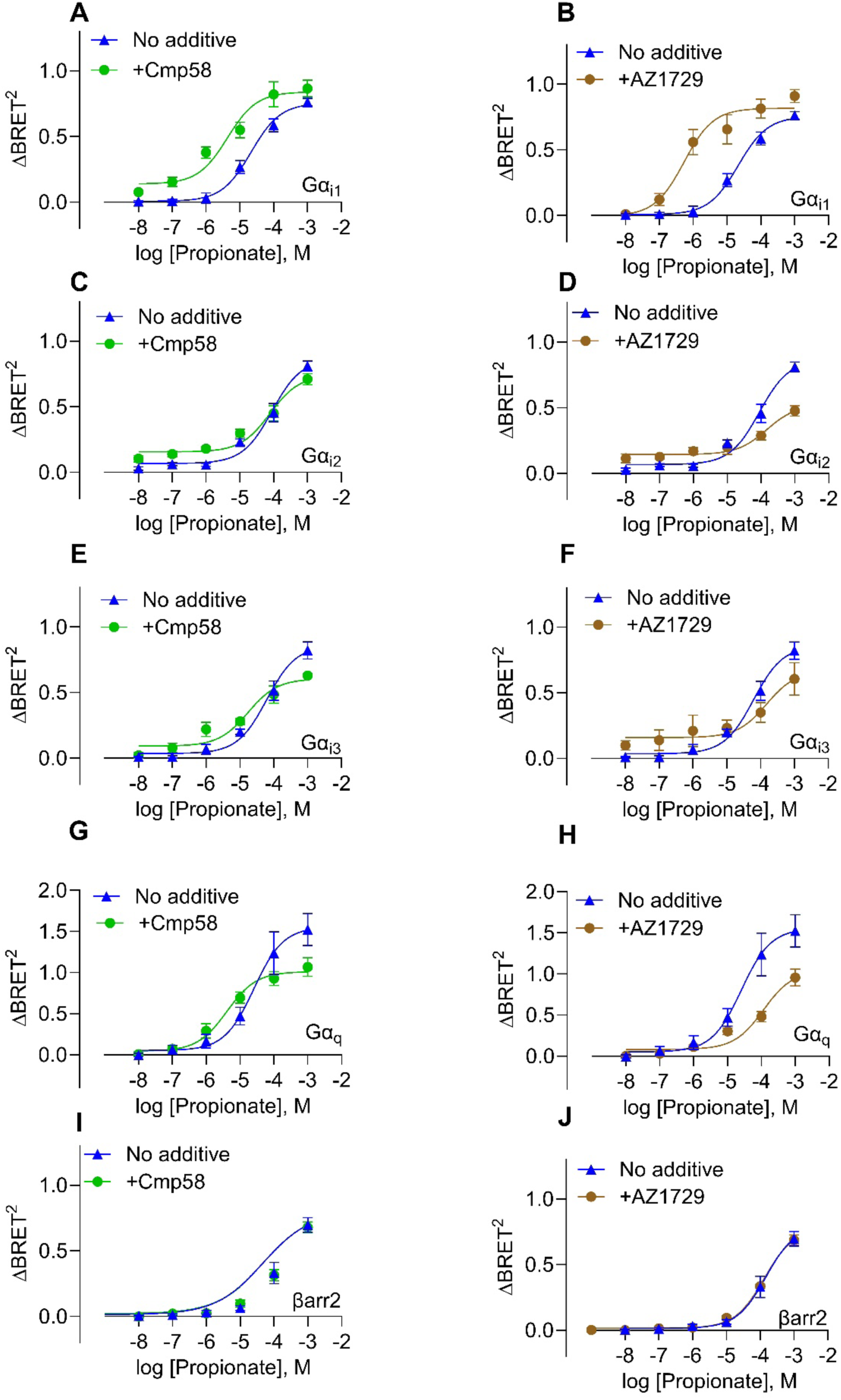
Positive allosteric effect of propionate is only evident through one G protein and allosteric binding do not affect the recruitment of β-arrestins. Dose–response curves for propionate (blue) in the absence or presence of Cmp58 (1 µM, 5-min preincubation; green) or AZ1729 (1 µM, 5-min preincubation; brown). Propionate-induced activation was measured for Gα_i1_ (**A**–**B**), Gα_i2_ (**C–D),** Gα_i3_ (**E–F**), Gα_q_ (**G–H**), and recruitment of β-arrestin2 (**I–J**). (**A-J**) subfigures present nonlinear fit dose concentration of the compound in log (molar) on the x-axis and the unitless ΔBRET^2^ signal on the y-axis of at least three biological replicates (n=3). Curves show propionate’s net allosteric effect after subtraction of vehicle and modulator-alone signals (Methods). All results in Figure 3 are from at least three biological replicates (n=3) performed in duplicates.

**Table 3:** EC_50_ values and max responses observed at concentration tested (ΔBRET^2^) for propionate with or without Cmp58 or AZ1729 using eBRET. “–“ indicate no activity in that pathway. **\***Note that Cmp58 and AZ1729 alone for Gα_15_ activation were too potent for use at the higher concentration, and a lower concentration was therefore chosen.

| ebBRET<br>measurement | Propionate |  | +Cmp58 |  | +AZ1729 |  |
| --- | --- | --- | --- | --- | --- | --- |
|  | EC<sub>50</sub> (μM) | Max<br>response | EC<sub>50</sub><br>(μM) | Max<br>response | EC<sub>50</sub> (μM) | Max<br>response |
| Gα<sub>i1</sub> | 21.4 | 0.7552 | 4.265 | 0.8407 | 0.5339 | 0.8173 |
| Gα<sub>i2</sub> | 88.98 | 0.8652 | 77.84 | 0.7404 | 139.3 | 0.481 |
| Gα<sub>i3</sub> | 61.89 | 0.8587 | 16.57 | 0.485 | 169.2 | 0.5332 |
| Gα<sub>q</sub> | 25.56 | 1.554 | 4.078 | 1.01 | 115.5 | 1.043 |
| Gα<sub>11</sub> | 25.9 | 1.82 | 18.44 | 1.41 | 119 | 1.36 |
| Gα<sub>12</sub> | - | - | - | - | - | - |
| Gα<sub>15</sub><sup>*</sup> | 140.3 | 0.68 | 120 | 0.503 | 119 | 0.533 |
| Gα<sub>oB</sub> | 81.8 | 1.812 | 45.98 | 1.479 | 20.22 | 1.621 |
| βarr2 | 147.5 | 0.7987 | 148.9 | 0.7742 | 131.6 | 0.7754 |

In contrast to their positive effects on Gα_i1_, Cmp58 and AZ1729 reduced propionate-evoked activation of Gα_q_ and Gα_11_ (**Fig. 3G–H**, Supplementary Fig.S2G-J, **Table 3**). Both modulators lowered the maximal ΔBRET^₂^ signal for these pathways, and AZ1729 also shifted the propionate concentration–response curve to be less potent. Cmp58 had a minor effect on potency but still attenuated overall response magnitude, indicating that both ligands down-modulate Gα_q/11_-mediated signaling (Supplementary Fig.S2G-J, **Table 3**). The addition of either Cmp58 or AZ1729 did not have any effect on propionate-induced β-arrestin recruitment (**Fig. 3I-J**, Supplementary Fig.S2K-L).

Together, these results establish that Cmp58 and AZ1729 act as functionally selective allosteric FFA2R ligands, exhibiting dual modulatory behavior: potentiation of FFA2R recruitment of Gα_i1_ and inhibition of the Gα_q/11_ recruitment to the propionate-activated receptor with minimal allosteric effect on β-arrestin recruitment. Notably, within the Gα_i/o_ family, only Gα_i1_ was strongly enhanced by both modulators, whereas Gαi_2_ and Gαi_3_ were modestly negatively affected; a selectivity that underscores the precision of FFA2R allosteric preference.

### Lack of cooperativity in HEK293 cells for the two ago-PAMs Cmp58 and AZ1729

We next assessed whether the combination of Cmp58 and AZ1729 produced cooperative effects on FFA2R signaling, *allosteric activation,* as discovered in neutrophils^8^. In contrast to the neutrophil setting, co-application of Cmp58 and AZ1729 did not result in additional increase in either potency or efficacy in terms of G protein activation or β-arrestin 2 recruitment in HEK293 cells (**Fig. 4A–F**, **Table 2**). The signaling profiles of the combined treatment largely overlapped with those of the individual compounds, with no evidence of synergistic or cooperative activity. We also verified the β-arrestin recruitment using a NanoBiT complementation assay and excluded any assay-specific effects. Propionate induced a robust, concentration-dependent β-arrestin2 response (Supplementary Fig.S3A), where the addition of Cmp58 did not significantly shift potency or efficacy (Supplementary Fig.S3B). Cmp58, AZ1729, or their combination alone further failed to recruit β-arrestin2 above baseline (Supplementary Fig. S3C). These results confirm that while propionate efficiently engages β-arrestin2, the allosteric modulators primarily influence G protein coupling rather than β-arrestin recruitment. Thus, although both Cmp58 and AZ1729 alone exhibited agonistic properties and distinct G protein preferences compared with propionate, their combination did not produce cooperative *allosteric activation* under these conditions in HEK293 cells.

**Fig.4:**
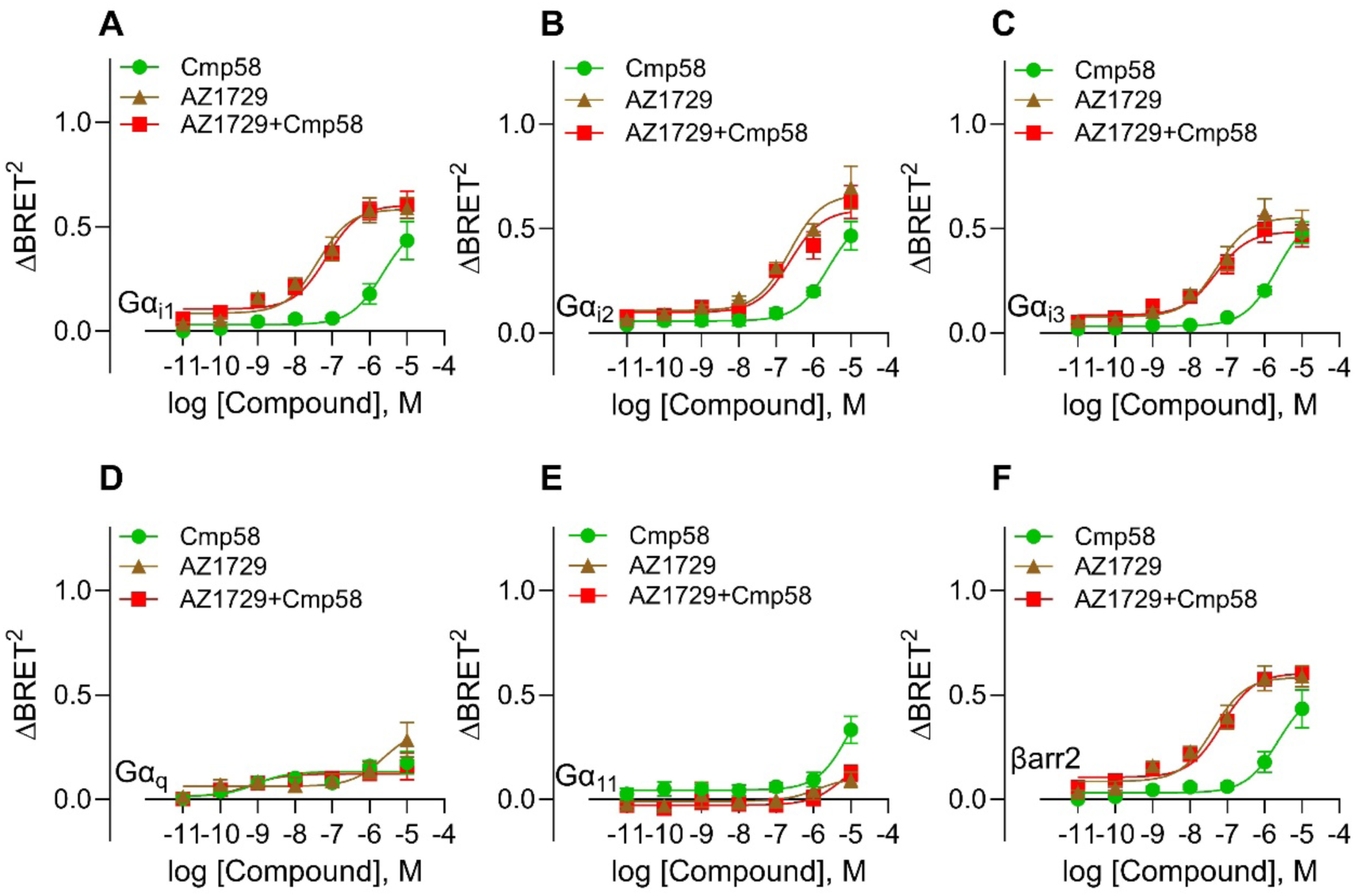
AZ1729 and Cmp58 do not cooperatively affect the G protein signaling and β-arrestin 2 recruitment. Concentration–response curves for the FFA2R allosteric agonists AZ1729 (brown), Cmp58 (green), and their combination (red) monitoring activation of (**A**) Gα_i1_, (**B**) Gα_i2_, (**C**) Gα_i3_, (**D**) Gα_q_, (**E**) Gα_11_, and (**F**) β-arrestin2 recruitment. (**A-F**) subfigures present nonlinear fit dose concentration of the compound in log (molar) on the x-axis and the unitless ΔBRET^2^ signal on the y-axis of at least three biological replicates performed in duplicates (n=3).

### Signaling and functional outcome of FFA2R activation in primary neutrophils

FFA2R signaling in neutrophils is well established and regulates both intracellular Ca^2+^ mobilization and NADPH oxidase–dependent generation of reactive oxygen species (ROS). These responses are triggered by the endogenous ligand propionate and can be further modulated allosterically by ligands such as Cmp58 and AZ1729^8,32^. ROS generated by NADPH oxidase are essential for neutrophil antimicrobial defense and act as secondary messengers in immune signaling pathways^34^. To provide a comparative reference for subsequent analyses in HL60 and HEK293 cells, we first quantified ROS and Ca^2+^ responses in primary human neutrophils under matched experimental conditions (temperature, buffer composition, ligand concentration ranges and addition sequence, and acquisition settings).

ROS production was first investigated in human neutrophils. Stimulation with propionate induced a rapid release of superoxide anions, which only occurs when cells are pre-stimulated with either allosteric modulator Cmp58 or AZ1729 (**Fig. 5A**). Neither modulator alone triggered detectable ROS formation, but co-application of Cmp58 and AZ1729 without propionate addition induced a strong response, an activation profile called *allosteric activation* (**Fig. 5B**).

**Fig.5:**
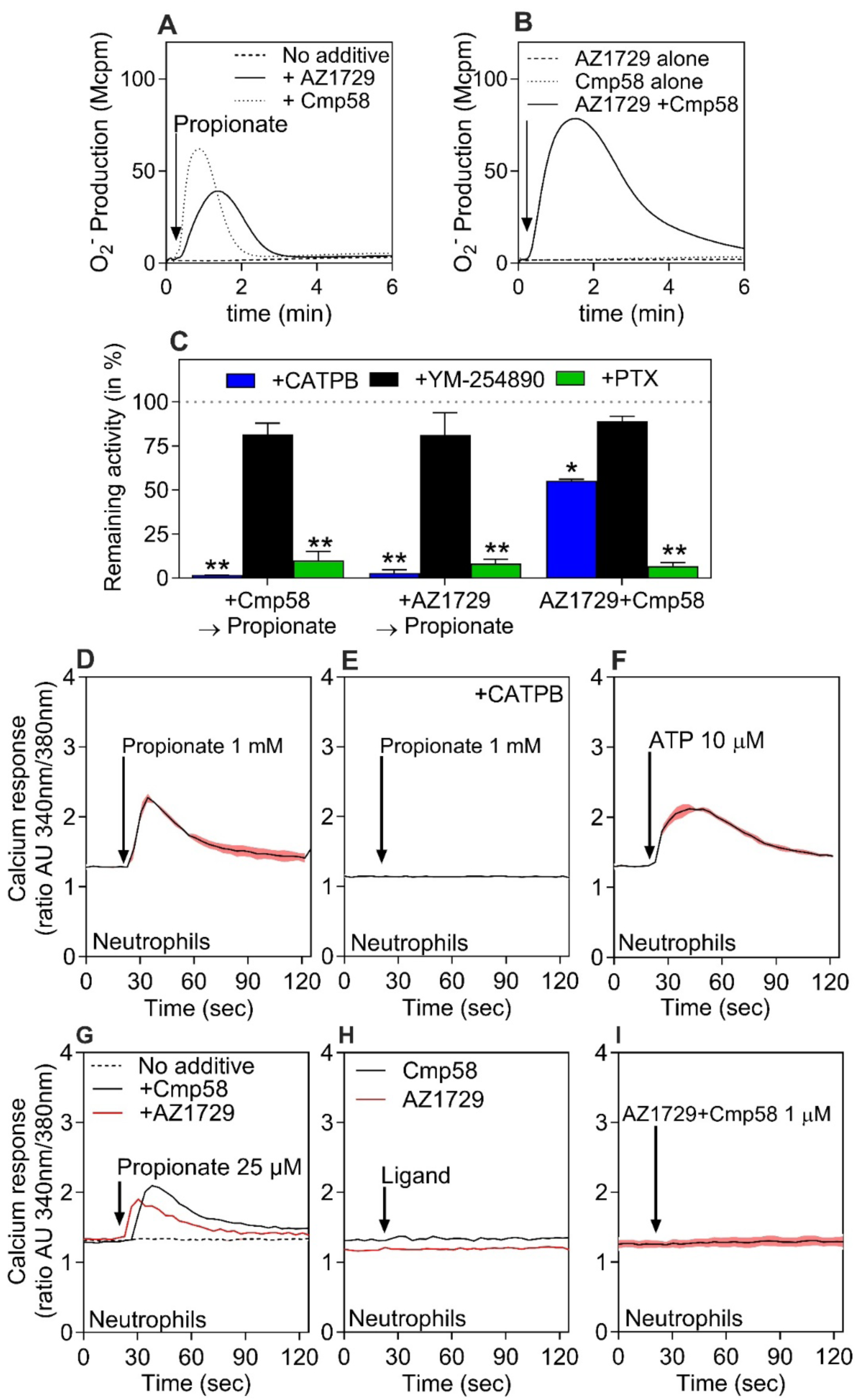
FFA2R-mediated superoxide generation and Ca^2+^ signaling in human neutrophils. (**A**) Neutrophils were stimulated with propionate (25 µM) alone (dashed line) or after 5-min preincubation with AZ1729 (1 µM, solid line) or Cmp58 (1 µM, dotted line). (**B**) Neutrophils were stimulated with AZ1729 (1 µM, dashed line), Cmp58 (1 µM, dotted line), or their combination (1 µM each, solid line). Superoxide production was monitored continuously and expressed as Mega counts per minute (Mcpm); traces are representative of five independent experiments. **(C)** O₂⁻ production induced by propionate in neutrophils sensitized with Cmp58, AZ1729, or AZ1729+Cmp58 in the presence of CATPB (FFA2R antagonist, 100 nM, 5 min; blue), YM-254890 (200 nM, 5 min; black), or pertussis toxin (500 ng/mL, 120 min; green). Data are shown as remaining activity (% of peak response without inhibitor; mean ± SEM, n = 3). Peak responses with and without inhibitor were compared using paired Student’s *t*-test. **(D–I)** Transient changes in intracellular calcium concentration ([Ca^2+^]_i_) in Fura-2–loaded neutrophils. Cells were stimulated with propionate (1 mM) (**D**), propionate (1 mM) after 5-min preincubation with CATPB (100 nM) (**E**), ATP (10 µM) (**F**), propionate (25 µM) in the absence (dotted line) or after 5-min preincubation with Cmp58 (1 µM, black) or AZ1729 (1 µM, red) (**G**), Cmp58 (1 µM, black) or AZ1729 (1 µM, red) alone (**H**), or the combination of AZ1729 and Cmp58 (1 µM each) (**I**). Ligand additions are indicated by arrows. Ca^2+^ traces are representative of three independent experiments; colored outlines denote SEM. Abscissa: time (s); ordinate: change in [Ca^2+^]_i_ expressed as the Fura-2 340/380-nm fluorescence ratio (arbitrary units).

To confirm the signaling profiles observed, pharmacological inhibitors were applied. The FFA2R antagonist CATPB fully abolished the allosterically modulated propionate-induced ROS release, while the Gα_q_ inhibitor YM-254890 had no significant effect on the response. Pertussis toxin (PTX) significantly reduced the response (**Fig. 5C**), suggesting that Gα_i/o_ proteins transduce the FFA2R-mediated signals that activate the NADPH oxidase.

One of the earliest events in GPCR activation is the increase of intracellular calcium in the cytosol ([Ca^2+^]_i_) following the addition of agonist stimulation^27,35^. Neutrophils also displayed a transient increase in intracellular Ca^2+^ following stimulation with propionate, which was blocked by CATPB, confirming a receptor-specific response (**Fig. 5D–E**). As a positive control, ATP, acting through the endogenous P2Y₂ receptor, induced a robust Ca^2+^ signal and served as a positive control (**Fig. 5F**). At lower propionate concentrations, Ca^2+^ responses were lower but could be allosterically enhanced by the addition of Cmp58 or AZ1729 (**Fig. 5G**). Neither Cmp58 nor AZ1729 alone, nor their combination, elicited Ca^2+^ release, which demonstrates that allosteric activation is *a functional selective* event (**Fig. 5H–I**).

Together, these findings illustrate the functional selectivity of FFA2R signaling in neutrophils. The endogenous ligand propionate primarily triggers transient Ca^2+^ mobilization without activation of the NADPH oxidase, whereas the allosteric modulators Cmp58 and AZ1729 act as true allosteric modulators, enhancing or enabling receptor activity only in the presence of propionate. When both modulators are combined in the absence of propionate, they induce ROS generation without transient Ca^2+^ mobilization, i.e. *allosteric activation*. Collectively, the results demonstrate that FFA2R engages distinct effector programs under different ligand contexts. To explore these allosteric mechanisms under more controlled and genetically accessible conditions, and to directly compare the functional outcomes across cell types, subsequent experiments were performed in the neutrophil-like HL60 model.

### Transient expression of FFA2R in HL60 cells reconstitutes ROS and Ca^2+^ signaling

The differentiated HL60 cells express the ROS generating NADPH oxidase that is activated by the phorbol ester PMA and the FPR1 agonist fMLF^36^. Expression of FFA2R did not alter responses to PMA or fMLF, confirming that the receptor-independent (i.e., through PMA) and receptor-dependent (i.e., through fMLF) ROS activation pathways remain intact in these cells (**Fig. 6A**).

**Fig.6:**
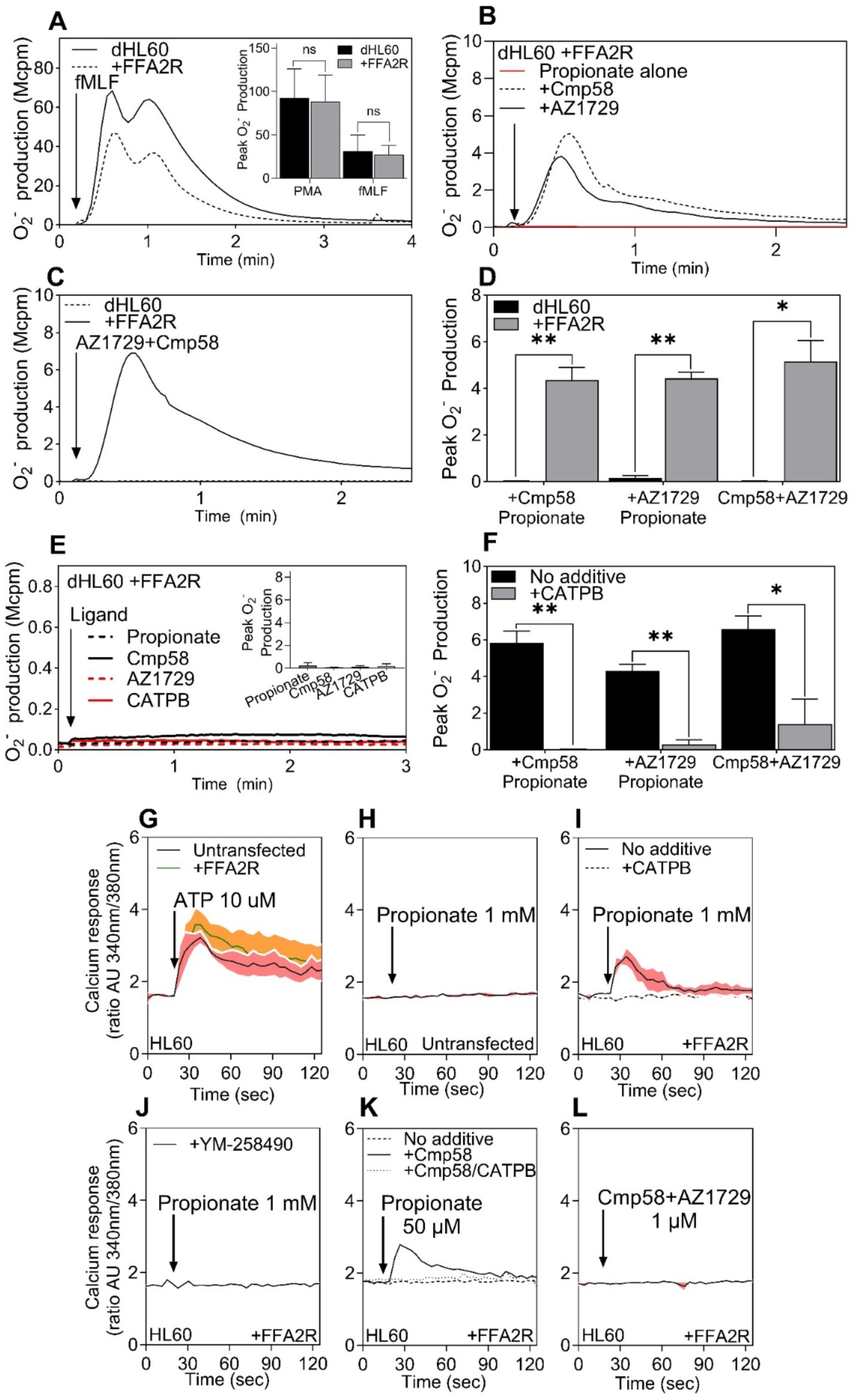
Transient expression of FFA2R in HL60 mimic the FFA2R signaling profile of neutrophils. **(A)** dHL60 (solid line) and dHL60+FFA2R (dashed line) cells were stimulated with fMLF (100 nM) after 5 min at 37 °C. *Inset:* quantification of peak O₂⁻ responses to fMLF (100 nM) and PMA (50 nM) in dHL60 (black) and dHL60+FFA2R (grey) cells (mean ± SEM, n = 3). “ns” indicates no significant difference between the two cell types (paired Student’s *t*-test). **(B)** dHL60+FFA2R cells stimulated with propionate (25 µM) alone or after 5-min preincubation with Cmp58 or AZ1729 (1 µM each). **(C)** dHL60 and dHL60+FFA2R cells stimulated with the combination Cmp58+AZ1729 (1 µM each). **(D)** Summary of peak O₂⁻ production in dHL60 (grey bar) and dHL60+FFA2R (black bar) for propionate in the presence of Cmp58 or AZ1729 and for Cmp58+AZ1729 (mean ± SEM, n = 3; paired Student’s *t*-test). **(E)** O₂⁻ production in dHL60+FFA2R in response to propionate (25 µM), Cmp58 (1 µM), AZ1729 (1 µM), or CATPB (1 µM); inset shows peak values (mean ± SEM, n = 3). **(F)** Peak O₂⁻ production induced by propionate (25 µM) in dHL60+FFA2R sensitized with Cmp58, AZ1729, or Cmp58+AZ1729 (1 µM) in the absence (black bar) or presence of CATPB (100 nM, 5 min, grey bar) (mean ± SEM, n = 3; paired *t*-test). **(G–L)** Transient changes in [Ca^2+^]_i_ in HL60 cells ± FFA2R expression. **(G)** ATP (10 µM) in untransfected and FFA2R-expressing HL60. **(H)** Propionate (1 mM) in untransfected HL60. **(I)** Propionate (1 mM) in FFA2R-expressing HL60 with (solid line) or without CATPB (100 nM, 5 min, dotted line). **(J)** Propionate (1 mM) in FFA2R-expressing HL60 preincubated with YM-258490 (200 nM, 5 min). **(K)** Propionate (50 µM, dashed line) in FFA2R-expressing HL60 with Cmp58 (1 µM, 5 min, solid line) and/or CATPB (1 µM, 5 min, dotted line). **(L)** Cmp58+AZ1729 (1 µM each) in FFA2R-expressing HL60. Ligand additions are indicated by arrows. Traces are representative of three independent experiments; shaded areas indicate SEM. Abscissa: time (s); ordinates: O₂⁻ production (Mcpm) or [Ca^2+^]_i_ as the Fura-2 340/380 ratio (arbitrary units).

FFA2R expression enabled robust ROS production in response to propionate, which similarly to neutrophils, only is activated in the presence by Cmp58 or AZ1729 (**Fig. 6B**). Co-application of Cmp58 and AZ1729 induced strong allosteric activation (**Fig. 6C**). No FFA2R activation was observed in untransfected dHL60 cells, underscoring a receptor-specific response (**Fig. 6D**). High concentrations of propionate or either Cmp58 or AZ1729 alone failed to trigger ROS production (**Fig. 6E**) and FFA2R responses could be fully blocked by the addition of FFA2R antagonist CATPB (**Fig. 6F**).

Although undifferentiated promyelocytic HL60 cells lack a functional NADPH oxidase, ATP stimulation of endogenous P2Y₂R elicited a clear Ca^2+^ transient (**Fig. 6G**), validating the Ca^2+^ assay in this background^28,37^. These cells could, thus, be used to determine the activity induced through this pathway by the FFA2R selective ligands. The response induced by ATP was not affected when FFA2R was transiently expressed in HL60 cells (**Fig. 6G**) and untransfected HL60 did not respond to propionate, even at high concentrations (**Fig. 6H**). FFA2R-expressing HL60 cells displayed clear Ca^2+^ transients upon propionate stimulation at high ligand concentrations, which could be abolished upon application of antagonist CATPB (**Fig. 6I)** or the Gα_q_ inhibitor YM-254890 (**Fig. 6J**). Positive allosteric modulation by Cmp58 was observed for propionate-induced response at lower concentrations of the agonist and this response was also inhibited by CATPB (**Fig. 6K**). The combination of Cmp58 and AZ1729, did not elicit Ca^2+^ release (**Fig. 6L**).

Taken together, transiently expressing FFA2R in HL60 cells re-established propionate-evoked Ca^2+^ mobilization and, upon DMSO differentiation, permitted allosterically driven ROS production, paralleling the responses seen in primary neutrophils.

### Transient expression of FFA2R in HEK293 cells reveals Gα_q_-dependent Ca^2+^ signaling

As Ca^2+^ mobilization represents a central signaling output of FFA2R in both neutrophils and HL60 cells, we next investigated whether transiently expressed FFA2R in HEK293 cells could induce Ca^2+^ mobilization responses. Propionate induced a clear concentration-dependent rise in intracellular Ca^2+^ concentration in FFA2R-expressing HEK293 cells, with strong responses observed at 1 mM and 500 µM (**Fig. 7A**). The signal was abolished by the FFA2R antagonist CATPB (**Fig. 7B**) and by the Gα_q_ inhibitor YM-254890 (**Fig. 7C**), confirming receptor and Gα_q_ dependence. At lower concentrations of propionate (250 µM and 100 µM), Ca^2+^ responses were lower but were significantly enhanced at 250 µM by preincubation with the allosteric modulator Cmp58 (**Fig. 7D–E**, **G–H**). By contrast, AZ1729 did not potentiate Ca^2+^ signals under the same conditions (**Fig. 7F, I**). Neither Cmp58 nor AZ1729 alone, nor their combination, triggered detectable Ca^2+^ release in the absence of propionate (**Fig. 7J–L**). To verify assay performance, a control experiment was performed with CATPB alone (Supplementary Fig.S3D) and YM-254890 alone (Supplementary Fig.S3E), where neither elicited a Ca^2+^ response. To validate the assay, ATP stimulation of the Gα_q_-coupled P2Y₂ receptor transiently expressed in HEK293 cells evoked a robust, transient Ca^2+^ rise that was abolished by YM-254890 (Supplementary Fig. S3F), confirming that the response faithfully reports Gα_q_-mediated Ca^2+^ mobilization.

**Figure 7.**
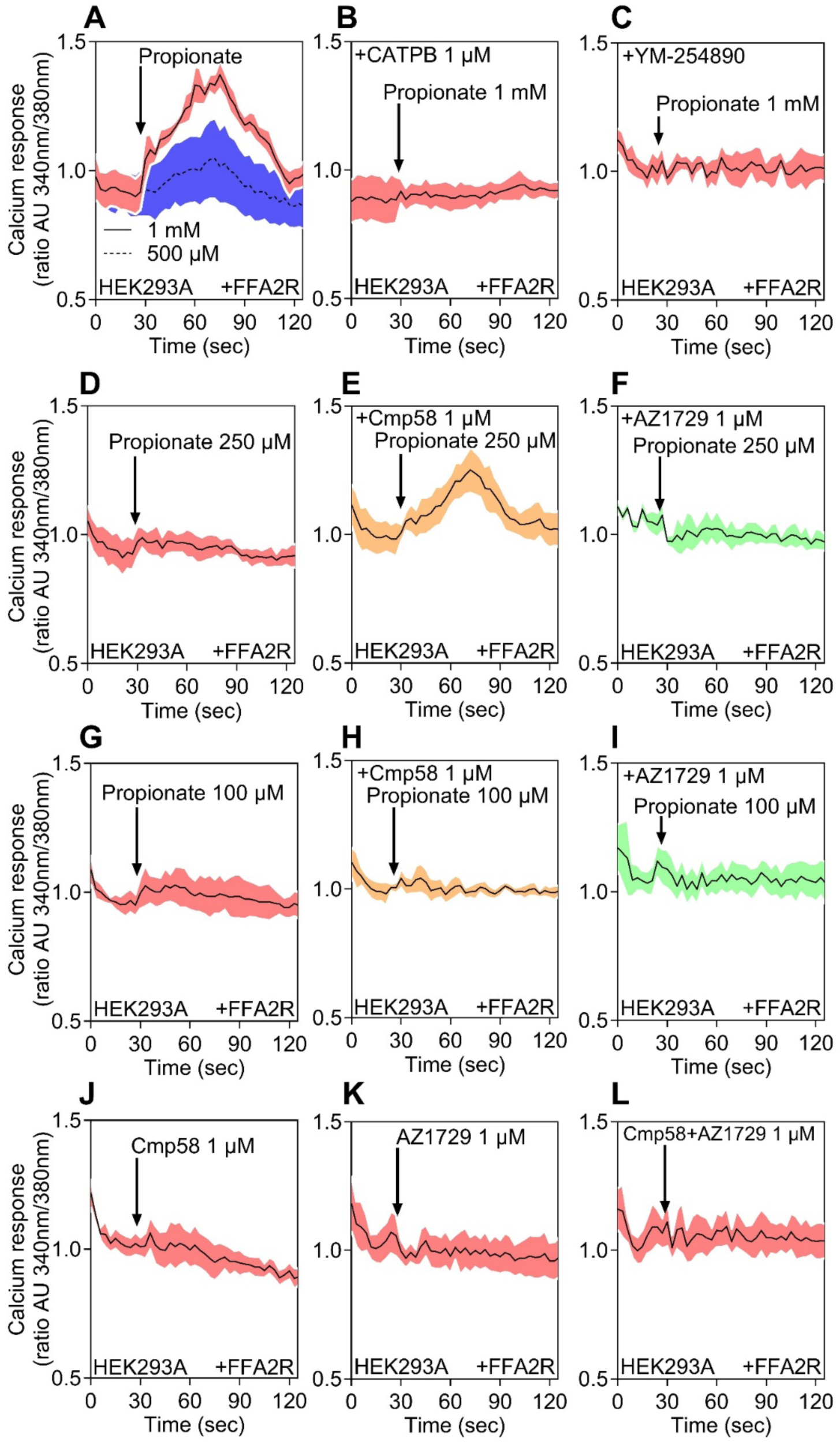
FFA2R-mediated Ca^2+^ mobilization in HEK293 cells transiently expressing FFA2R. (**A–L**) The transient rise of intracellular Ca^2+^ concentration was followed in HEK293T cells transiently expressing FFA2R as indicated. **(A**) Cells were stimulated with propionate (1 mM, solid line) or 500 μM (dashed line). **(B)** Cells were stimulated with propionate (1 mM) after 5 min preincubation with the FFA2R antagonist CATPB (1 μM). **(C)** Cells were stimulated with propionate (1 mM) after 5 min preincubation with the Gα_q_ inhibitor YM-254890 (200 nM). **(D)** Cells were stimulated with propionate (250 μM), or with propionate (250 μM) in the presence of the FFA2R allosteric modulators **(E)** Cmp58 (1 μM) or **(F)** AZ1729 (1 μM), each added 5 min before agonist. **(G)** Cells were stimulated with propionate (100 μM), or with propionate (100 μM) in the presence of **(H)** Cmp58 (1 μM) or **(I)** AZ1729 (1 μM). **(J–L)** Cells were stimulated with Cmp58 (1 μM), AZ1729 (1 μM), or the combination of both modulators (1 μM each). Ligand addition is indicated by arrows. Data are representative of three independent experiments, and shaded areas represent the standard error of the mean. Abscissa, time of study (sec); ordinate, increase in intracellular Ca^2+^ [Ca^2+^]_i_ given as the change in the ratio between Fura-2 fluorescence at 340 and 380 nm (AU, arbitrary units).

Taken together, these data showed that FFA2R coupled to Gα_q_-dependent Ca^2+^ mobilization in HEK293 cells, with Cmp58 selectively enhancing propionate responses. Although Cmp58 and AZ1729 activated Gα_q_ in ebBRET assays, neither compound alone elicited Ca^2+^ release in HEK293, neutrophils, or HL60 cells. Viewed alongside neutrophil and HL60 datasets, the results were consistent with a conserved FFA2R-driven Ca^2+^ response across these systems.

## Discussion

FFA2R provides a tractable system to examine how allosteric ligands shape GPCR signaling in distinct cellular milieus. Across HEK293, HL60, and primary neutrophils, our data reveal pathway-selective modulation by Cmp58 and AZ1729 and a cell-type dependence for downstream effector engagement. We therefore interpret the findings around three themes: (i) selective enhancement or dampening of defined G protein pathways, (ii) emergence of myeloid-restricted oxidative responses and cooperative allosteric activation, and (iii) alignment with recently reported dual-site architecture of FFA2R, offering a mechanistic context for the observed ligand interactions. This framework explains why identical ligands yield divergent outputs across cells while preserving a conserved Ca^2+^ signal.

### FFA2R signaling profile in HEK293 cells

Using ebBRET in HEK293 cells, we found that the endogenous ligand propionate activates most G proteins, with the exception of Gα_12_. In contrast, the allosteric agonists Cmp58 and AZ1729 behaved as biased ligands, activating a more restricted set of G protein pathways and inducing little to no β-arrestin recruitment. Propionate-induced dose-dependent recruitment of both β-arrestin1 and 2 consistent with previous reports^38^. Prior studies suggested FFA2R coupling to Gα_i/o_ and Gα_q_^25,39,40^, and yeast chimeric systems hinted at possible coupling to Gα ^39^ but our results now validate G protein coupling profiles in a mammalian system. AZ1729 has been reported as a selective Gα_i_ agonist with positive allosteric effects on Gα_i_ and negative modulation of Gα_q_^7^, while Cmp58 inhibits adipocyte lipolysis *via* Gα_i_ in a PTX-sensitive manner^19^. In HEK293 cells, ebBRET showed pathway-specific bidirectional modulation by Cmp58 and AZ1729: Gα_i1_ was potentiated, Gα_i2_ and Gα_i3_ were attenuated, Gα_q/11_ was reduced, and β-arrestin2 recruitment was unchanged.

### Shared mechanisms of FFA2R allosteric modulation and activation in neutrophils and HL60 cells

FFA2R represents a unique GPCR model for dissecting allosteric modulation because of its multiple allosteric binding sites and the ability of distinct ligands to produce highly specific signaling outcomes^8,11,13,20^. In primary human neutrophils, FFA2R activation by the endogenous short-chain fatty acid propionate elicited transient Ca^2+^ mobilization but did not stimulate NADPH oxidase–dependent ROS production. Cmp58 and AZ1729 act as true allosteric modulators in primary human neutrophils, allowing propionate to drive ROS production, while also amplifying Ca^2+^ mobilization. These modulators therefore expand FFA2R signaling capacity by coupling propionate to both oxidative and Ca^2+^-dependent responses. A particularly striking feature of FFA2R pharmacology is the phenomenon of allosteric activation, observed when Cmp58 and AZ1729 are applied in combination in lieu of an orthosteric ligand. Under these conditions, the two ligands cooperatively induce ROS without eliciting Ca^2+^ mobilization, responses that neither ligand alone can trigger.

To further explore these mechanisms in a genetically tractable system, FFA2R was transiently expressed in HL60 cells. In this model, FFA2R expression restored responsiveness to propionate, leading to Ca^2+^ mobilization and allosterically modulated ROS generation. As in neutrophils, co-application of Cmp58 and AZ1729 induced allosteric activation, whereas neither ligand alone could trigger activation. All responses were fully abolished by CATPB and were absent in untransfected cells, confirming a receptor dependent activation. Together, these findings demonstrate that HL60 cells mimic the key features of neutrophil FFA2R signaling, including both allosteric modulation and allosteric activation. The ability to manipulate receptor expression and experimental conditions makes HL60 cells a powerful and complementary model for detailed mechanistic studies of FFA2R signaling and may also be adapted to investigate the function of other GPCRs under controlled and reproducible experimental conditions.

### Allosteric activation of FFA2R is context-dependent and shared between neutrophils and HL60 cells

Allosteric activation is an emerging concept in GPCR pharmacology, describing a scenario where two allosteric modulators positively modulate each other by binding to distinct sites on the same receptor without involvement of an orthosteric ligand^8,20^. In neutrophils, the combination of Cmp58 and AZ1729 trigger ROS production without a Ca^2+^ signal, showing that type of activation is also functional selective^8^. A similar pattern is observed in HL60 cells, where co-application of the two modulators produces strong allosteric activation of the NADPH oxidase, again without measurable Ca^2+^ mobilization.

The consistent responses across neutrophils and HL60 cells indicate that allosteric activation relies on shared downstream components that couple FFA2R–Gα_i/o_ signaling to the NADPH oxidase system. This conclusion is supported by the sensitivity of the response to pertussis toxin, which abolishes ROS formation, while Gα_q_ inhibition has no effect on ROS production and the two allosteric modulators cooperatively do not affect Ca^2+^ signaling. The dissociation between Ca^2+^ and ROS responses suggests that the cooperative effect of Cmp58 and AZ1729 occurs downstream of the G protein, at a signaling node linking Gα_i/o_ activation to NADPH oxidase regulation.

In contrast, no allosteric activation was detected in HEK293 cells, where the combination of Cmp58 and AZ1729 failed to synergize to produce a stronger G protein or β-arrestin activation profile. Instead, both compounds functioned alone as strong agonists in this system, directly activating multiple G proteins without evidence of interdependent modulation. The absence of allosteric activation in HEK293 cells implies that this phenomenon requires cellular components present in myeloid cells but lacking in HEK293 cells. Interestingly, however, their strongest positive modulation in HEK293 was observed for Gα_i1_, aligning with the pertussis toxin sensitivity seen in neutrophils. This convergence suggests that allosteric activation is at least partly mediated through Gα_i/o_-dependent mechanisms. Together, these findings demonstrate that FFA2R allosteric activation is a cell type–dependent event, emerging only in contexts where FFA2R signaling converges on other downstream pathways. This highlights how receptor allosterism can be shaped not only by ligand-receptor interactions but also by the signaling architecture of the cell in which the receptor resides.

### Ca^2+^ signaling across all three systems

In addition to G protein profiling, we provide the first systematic characterization of FFA2R-mediated Ca^2+^ signaling in HEK293 cells using Fura-2. Propionate triggered robust Ca^2+^ transients that were abolished by the FFA2R antagonist CATPB and the Gα_q_ inhibitor YM-254890, confirming receptor and Gα_q_ dependence. Cmp58 enhanced Ca^2+^ responses at submaximal propionate concentrations, whereas AZ1729 had no effect, and neither modulator triggered Ca^2+^ release alone. These results establish that FFA2R couples efficiently to Gα_q_-mediated Ca^2+^ signaling in HEK293 cells and that Cmp58 selectively enhances this pathway. Importantly, Ca^2+^ signaling provides a functional endpoint that can be directly compared across cell systems. When viewed alongside the data from neutrophils and HL60 cells, Ca^2+^ mobilization emerges as a conserved functional response across all examined systems. This is consistent with the role of Ca^2+^ signaling as one of the earliest activation events downstream of GPCR stimulation, acting as a rapid and sensitive indicator of receptor engagement. While in myeloid cells FFA2R signaling extends to NADPH oxidase–dependent ROS generation, the Ca^2+^ response remains a consistent and receptor-proximal event. The selective enhancement of Ca^2+^ signaling by Cmp58 in all systems further indicates that this pathway represents a shared point of allosteric control, even though the downstream consequences of receptor activation diverge according to cellular context. Notably, allosteric activation by Cmp58 and AZ1729 induce ROS generation which occurs independently of Ca^2+^ mobilization, suggesting that additional, yet undefined, signaling intermediates act downstream of Gα_i/o_ for FFA2R to activate the oxidative burst.

### Structural considerations and multiple ligand binding sites

Structural studies have revealed at least three binding sites in FFA2R, including two distinct allosteric pockets^41^. The orthosteric site is shallow, and molecular dynamics simulations suggest transient interactions between propionate and surrounding residues. The allosteric agonist 4-CMTB binds one of the allosteric sites, stabilizing the active receptor conformation and enhancing orthosteric agonist potency^41^. Our previous work showed that 4-CMTB had weaker priming effects than Cmp58 but could still activate neutrophil NADPH oxidase in combination with propionate^8^. Furthermore, 4-CMTB could activate neutrophils with AZ1729, but not with Cmp58, suggesting that Cmp58 and 4-CMTB share the same binding site, whereas AZ1729 binds the second allosteric site^8^.

Recently, cryo-EM structures of FFA2R resolved the two distinct allosteric sites: site 1, near TM6, where 4-CMTB (same site as Cmp58) binds, and site 2, above ICL2, where AZ1729 interacts to stabilize a Gα_i/o_-favored conformation^11^. These structural insights align closely with our functional data, showing that Cmp58 enhances Gα_q_-mediated Ca^2+^ signaling, while AZ1729 promotes Gα_i/o_-dependent ROS production. When combined, the ligands cooperatively induce *allosteric activation*, a Ca^2+^-independent, pertussis-toxin–sensitive response consistent with joint engagement of both sites. Thus, structural and functional evidence together support a dual-site mechanism in which site 1 drives TM6-mediated Gα_q_ activation and site 2 modulates ICL2 to bias FFA2R signaling toward Gα_i/o_ pathways, explaining the context-dependent receptor behavior observed across our model systems.

### Physiological context for FFA2R-mediated signaling

FFA2R is a useful model for studying GPCR allosterism, with a broad pharmacological toolkit that includes orthosteric agonists, antagonists, and several allosteric modulators with distinct signaling properties^7,21,42,43^. Prior work has begun to define the molecular basis of its allosteric control, suggesting that FFA2R can support both positive modulation and cooperative interactions between ligands^7,8,13,17,19,44^.

Beyond its mechanistic appeal, FFA2R has clear physiological relevance: it mediates host responses to short-chain fatty acids (SCFAs) derived from microbial fermentation and contributes to the regulation of energy metabolism, inflammation, and immune defense^16,45–48^. In mouse models, FFA2R has been linked to SCFA-associated pathophysiology, including colitis, asthma, and arthritis^48–50^. Consistent with this, small-molecule FFA2R agonists and antagonists are being developed for inflammatory and metabolic diseases, including colitis, dermatitis, asthma, and obesity-related disorders^51,52^

SCFAs exert protective effects in inflammatory bowel disease through FFA2R^48^, while FFA2R deficiency promotes obesity even under normal dietary conditions and overexpression in adipose tissue protects against diet-induced obesity^53^. Reports that some allosteric ligands, such as 4-CMTB (that bind to the same allosteric site site as Cmp58^8^), enhance insulin secretion and β-cell function further underscore potential translational interest, while highlighting the need for careful validation across systems^54^. Our findings on FFA2R allosterism outline how ligands might be optimized for specific immune and metabolic responses.

Taken together, these data separate conserved from context-dependent outputs of FFA2R activation. Ca^2+^ mobilization is a shared, receptor-proximal readout that Cmp58 consistently potentiates, whereas ROS generation is confined to myeloid settings and depends on Gα_i/o_-linked machinery that is absent in HEK293 cells. Importantly, Cmp58 and AZ1729 produce allosteric activation in neutrophils and HL60, yielding robust ROS without a Ca^2+^ transient; this cooperative, pertussis toxin sensitive effect is not seen in HEK293, consistent with a downstream integration node. These differences likely reflect cell-specific components such as the NADPH oxidase complex and cytoskeletal regulation. Using three complementary models provides physiological relevance (neutrophils), experimental tractability (HL60), and reductionist control (HEK293), yielding a fuller view of FFA2R allosterism. This framework should guide hypothesis-driven development of cell-selective FFA2R modulators and help avoid over-generalization of signaling event across biological contexts.

### Conclusions

Across three human cell systems, we define how orthosteric and allosteric ligands influence FFA2R signaling in a context-dependent manner. In HEK293 cells, ebBRET profiling showed that propionate activates multiple G proteins, with the excpeption of Gα_12_, whereas Cmp58 and AZ1729 act as pathway-selective ago-modulators that enhance Gα_i1_, reduce Gα_q/11_, and have minor effect on β-arrestin2. Because ebBRET reports proximity under overexpression and therefore profiles coupling capacity rather than endogenous efficacy, we interpret these HEK293 data as pathway-availability maps, validated against downstream functional readouts. Functional assays then established a conserved role for FFA2R in Gα_q_-dependent Ca^2+^ mobilization: propionate evokes robust Ca^2+^ signals and Cmp58 selectively potentiates submaximal responses, while AZ1729 does not share this effect.

In myeloid systems, ligand outcomes diverge at the level of effector coupling. In primary neutrophils and FFA2R-transfected HL60 cells, propionate alone raises intracellular Ca^2+^ but requires either Cmp58 or AZ1729 to drive NADPH oxidase–dependent ROS. Interestingly, co-application of Cmp58 and AZ1729 produces allosteric activation, a pertussis-toxin–sensitive ROS burst that occurs without a Ca^2+^ transient, placing this cooperative process downstream of Gα_i/o_ and upstream of NADPH oxidase. Here, we use *allosteric activation* descriptively to denote cooperative responses of Cmp58 and AZ1729; however the precise molecular mechanism remains to be defined. These cell-specific outcomes are consistent with cryo-EM models resolving two discrete allosteric pockets on FFA2R, one TM6-adjacent and Cmp58/4-CMTB preferring, the other ICL2-proximal and AZ1729 preferring, and co-occupancy provides a structural rationale for the cooperative, PTX-sensitive ROS response observed in myeloid cells.

Taken together, these data indicate that cellular architecture, more than receptor structure alone, gates whether FFA2R outputs terminate in Ca^2+^ mobilization or extend to an oxidative burst. The comparative use of HEK293, HL60, and neutrophil systems therefore supports a hierarchical model: propionate consistently drives receptor-proximal Ca^2+^ mobilization *via* Gα_q/11_, while Cmp58 and AZ1729 redirect signaling toward Gα_i/o_ and, in myeloid cells, cooperatively engage the oxidative burst.

Collectively, these results argue that a multi-assay framework spanning receptor-proximal coupling and downstream effectors is required to fully capture allosteric modulation at FFA2R. Cross-examining G-protein coupling, Ca^2+^ mobilization, β-arrestin recruitment, and specialized cellular functions provides a more comprehensive understanding of ligand behavior and a stronger foundation for designing modulators with the desired pathway selectivity in the cellular target of interest.

## MATERIALS AND METHODS

### Reagents

Penicillin/streptomycin, trypsin, fetal bovine serum, reagent reservoirs, Nunclon Delta surface 6-well plates, Nunc EasYFlask 75 cm² flasks, Lipofectamine 2000, RPMI 1640 (without phenol red, supplemented), Poly-D-Lysine,Dulbecco’s modified Eagle’s medium (DMEM), phosphate-buffered saline (PBS), Hanks’ balanced salt solution (HBSS), OPTI-MEM, salmon sperm DNA solution and Fura-2 AM were purchased from Thermo Fisher Scientific (Waltham, MA, USA). Dextran and Ficoll-Paque were obtained from GE Healthcare Bio-Science (Uppsala, Sweden). Horseradish peroxidase (HRP) was obtained from Boehringer Mannheim (Mannheim, Germany). Cmp58 ((S)-2-(4-chlorophenyl)-3,3-dimethyl-N-(5-phenylthiazol-2-yl)butanamide) and AZ1729 were obtained from TOCRIS (Bristol, UK). The Nano-Glo® Luciferase Assay System and the Promega GloSensor-22F cAMP kit were purchased from Promega (Madison, WI, USA). Coelenterazine 400a was purchased from Nanolight Technologies (Pinetop, AZ, USA). YM-254890 was purchased from Wako Chemicals (Neuss, Germany). Pertussis toxin (PTX), Phorbol 12-myristate 13-acetate (PMA), N-formyl-Met-Leu-Phe (fMLF), EGTA, dimethyl sulfoxide (DMSO), bovine serum albumin (BSA), isoluminol, CATPB, TNF-α, propionic acid, forskolin, and ATP were purchased from Sigma Chemical Co. (St. Louis, MO, USA).

The ligands were dissolved according to the manufacturers’ recommendations and stored at −80 °C until use. Subsequent dilutions of receptor ligands and other reagents were made in PBS when performing BRET experiments or in Krebs-Ringer glucose phosphate buffer (120 mM NaCl, 4.9 mM KCl, 1.7 mM KH₂PO₄, 8.3 mM Na₂HPO₄, 1.2 mM MgSO₄, 10 mM glucose, and 1 mM CaCl₂ in dH₂O, pH 7.3) when neutrophils or HL-60 cells were used.

### Plasmids and ebBRET biosensor constructs

BRET experiments were performed as described previously^31^. The following biosensor constructs were used: rGFP-CAAX^55^, Rap1GAP-RlucII^25^, p63RhoGEF-RlucII^25^, PDZRhoGEF-RlucII^25^., β-arrestin1-RlucII^56^, and β-arrestin2-RlucII^56^, each described previously. FFA2R (GPR43), Gα_i1_, Gα_i2_, Gα_i3_, Gα_oA_, Gα_oB_, Gα_z_, Gα_q_, Gα_11_, Gα_12_, Gα_13_, Gα_14_, and Gα_15_ were purchased from cDNA.org (Bloomsburg University, Bloomsburg, PA). GRK2 was generously provided by Dr. Antonio De Blasi (Instituto Neurologico Mediterraneo Neuromed, Pozzilli, Italy).

G protein activation was measured using selective effector–RlucII biosensors with rGFP-CAAX as a plasma membrane anchor, together with the indicated Gα subunit and human FFA2R.

- Gα_i/o_ family activation was monitored using Rap1GAP–RlucII and rGFP–CAAX along with Gα_i1_, Gα_i2_, Gα_i3_, Gα_oA_, Gα_oB_, or Gα_z_.
- Gαq/11 family activation was assessed using p63RhoGEF–RlucII and rGFP–CAAX with Gα_q_, Gα_11_, Gα_14_, or Gα_15_.
- Gα_12/13_ activation was evaluated using PDZ–RhoGEF–RlucII and rGFP–CAAX with either Gα_12_ or Gα_13_.

β-arrestin recruitment to the plasma membrane was determined using β-arrestin1–RlucII or β-arrestin2–RlucII with or without GRK2, co-expressed with rGFP–CAAX and FFA2R.

### Transfection of HEK293 cells

HEK293T cells (*RRID:CVCL_0063*) were grown in plastic flasks at 37 °C with 5% CO_2_ in DMEM supplemented with 10% fetal calf serum, streptomycin (0.1 mg/ml), and penicillin (100 U/mL). The cells were passaged every two days and to avoid overgrowth and were transfected around 60–80% confluency. Cells were regularly tested for mycoplasma contamination.

For ebBRET sensors, HEK293T cells were transfected with WT FFA2R, and G protein or *β*-arrestin as described previously^31^. To ensure that all wells contained the same amount of DNA, salmon sperm was used as empty vector. Cells were transfected with Lipofectamine2000 in accordance with the manufacture’s protocol and subsequently seeded onto Poly-D-Lysine precoated 96-well plates and incubated for 48 hours at 37 °C with 5% CO_2_.

For the NanoBiT assay, HEK293T cells were transfected with FFA2R-SmBiT and *β*-arrestin-LgBiT. To ensure that all wells contained the same amount of DNA, salmon sperm was used as empty vector. Cells were transfected with Lipofectamine2000 in accordance with the manufacture’s protocol and subsequently seeded onto Poly-D-Lysine precoated 96-well plates and incubated for 48 hours at 37 °C with 5% CO_2_.

For Ca^2+^ mobilization, HEK293T cells were transiently transfected with human FFA2R (500 ng/well) using Lipofectamine2000 according to the manufacturer’s instructions. To ensure equal total DNA amounts across wells, salmon sperm DNA was used as an empty vector control. After transfection, cells were seeded onto poly-D-lysine–coated 96-well plates and incubated for 48 h at 37 °C in 5% CO₂ before performing Ca^2+^ measurements.

### Ligand-induced BRET measurements

G protein activation and β-arrestin recruitment were monitored using the enhanced bystander bioluminescence resonance energy transfer (ebBRET) platform as described previously ^25,31^. Briefly, transfected cells were washed twice with Hank’s balanced salt solution (HBSS) and incubated with coelenterazine 400a (5 µM, 5 min, 37 °C). When two ligands were tested sequentially, the second ligand was added immediately after coelenterazine incubation. Following pre-incubation, BRET signals were recorded as baseline readings before and after ligand or vehicle addition to detect ligand-induced changes in energy transfer.

BRET measurements were acquired at 37 °C using a CLARIOstar plate reader (BMG Labtech) equipped with monochromators for BRET² detection at 410/80 nm (donor, RlucII) and 515/30 nm (acceptor, rGFP). Data were expressed as the BRET ratio (acceptor/donor emission) and analyzed as ligand-induced ΔBRET² relative to baseline.

### Measuring Luminescence using the Nano-Glo® Luciferase Assay System

The Nano-Glo® Luciferase Assay System was followed using the provided manufacturers protocol. Briefly, the cell growth media was washed twice and replaced with 90 µL HBSS. The Nano-Glo® Luciferase Assay Substrate was diluted in Nano-Glo® Luciferase Assay Buffer and 10 μL per well was added. After 2 min at 37 °C, the luminescence was measured in two consecutives followed by addition of ligand or vehicle control and subsequent luminescence reads to detect ligand-induced changes in luminescence.

### HL60 cultivation, transfection and differentiation

HL60 cells were cultured under sterile conditions at 37° C in 5% CO_2_ in RPMI 1640 medium supplemented with 10% fetal calf serum (FCS), 2 mM L-glutamine, 1 mM sodium pyruvate, 100 units/mL penicillin and 100 μg/mL streptomycin (RPMI-complete medium). Cell cultures were passaged three times per week, maintaining cell densities between 1 x 10^5^ cells/mL × – 1 × 10^6^ cells/mL in 75 cm^2^ Nunc Flasks. To differentiate HL60 cells, the cells were seeded at a density of 2 x 10^5^ cells/mL in a 6 well Nunc plate and differentiated towards a non-adherent neutrophil-like phenotype (dHL60) by incubation with RPMI 1640 complete media supplemented with 1.3% DMSO for five days. To perform a transient expression of FFA2R in HL60 cells, undifferentiated HL60 cells were transfected with 1000 ng FFA2R at 350 000 cells/mL (5 mL total) using Lipofectamine2000 in accordance with the manufacturer’s protocol and subsequently seeded onto 6 well plates for 48 hours. The FFA2R-expressing HL60 cells were then counted and reseeded to density of 2×10^5^ cells/mL and subsequently differentiated using RPMI complete supplemented with 1.3 % DMSO for 5 days. To prepare the cells for measurement of NADPH-oxidase activity, the cells were washed and re-suspended to 1×10^6^ cells/ml in KRG, stored on ice until use on day five after start of the differentiation.

### Isolation of human neutrophils

Human neutrophils were isolated from buffy coats by dextran sedimentation followed by Ficoll-Paque gradient centrifugation, as originally described by Bøyum ^57^. Residual erythrocytes were removed by hypotonic lysis, after which neutrophils were washed and resuspended in Krebs–Ringer glucose (KRG) buffer. The final cell suspensions contained >90% neutrophils with a viability exceeding 95%. To enhance receptor responsiveness, cells were primed with TNF-α (10 ng/mL, 20 min, 37 °C) before use. Neutrophils were kept on ice after priming until immediately prior to stimulation.

### Measuring NADPH oxidase activity in neutrophils and dHL60 cells

Superoxide generation, reflecting NADPH oxidase–dependent production of reactive oxygen species (ROS), was measured using the isoluminol-enhanced chemiluminescence (CL) assay as previously described^58^. Measurements were performed in a six-channel Biolumat LB 9505 luminometer (Berthold Co., Wildbad, Germany) using disposable 4-mL polypropylene tubes containing a 900-µL reaction mixture of 1 × 10⁵ neutrophils or dHL60 cells, isoluminol (0.2 µM), and horseradish peroxidase (HRP; 4 U/mL). Tubes were equilibrated for 5 min at 37 °C before addition of agonist (100 µL), after which light emission was recorded continuously. When testing the effects of receptor-specific antagonists or allosteric modulators, these compounds were added to the reaction mixture 5 min prior to stimulation. Control cells treated identically but without antagonist or modulator were run in parallel for comparison.

### Ca^2+^ mobilization assay in neutrophils and HL60 cells

Neutrophils and HL60 cells were resuspended in Ca^2+^-free KRG buffer supplemented with 0.1% BSA at a density of 5 × 10⁷ cells/mL for neutrophils and 5 × 10⁶ cells/mL for HL60 cells. Cells were loaded with Fura-2 AM (5 µM) for 30 min at room temperature in the dark, then diluted 1:1 in RPMI 1640 medium without phenol red, and centrifuged at 900 × g. The cell pellets were washed once with KRG and resuspended in the same buffer to a final density of 2 × 10⁷ cells/mL (neutrophils) or 2.5 × 10⁶ cells/mL (HL60 cells).

Ca^2+^ measurements were performed using a PerkinElmer LC50 fluorescence spectrophotometer (PerkinElmer, Waltham, MA, USA) with excitation wavelengths of 340 nm and 380 nm and an emission wavelength of 509 nm (slit widths 5 nm and 10 nm, respectively). The transient increase in intracellular Ca^2+^ concentration was expressed as the ratio of fluorescence intensities (F340/F380) recorded over time.

### Ca^2+^ mobilization in HEK293T cells

HEK293T cells were maintained in DMEM supplemented with 10% fetal bovine serum and 1% penicillin–streptomycin at 37 °C in 5% CO₂. For transient expression, cells at 40–80% confluence were transfected with 500 ng FFA2R plasmid DNA per well (6-well format) using Lipofectamine 2000 according to the manufacturer’s instructions; salmon sperm DNA was added as carrier to equalize total DNA across conditions. 48 hours after transfection, cells were detached, washed in Ca^2+^-free Krebs–Ringer glucose (KRG) buffer containing 0.1% BSA, and loaded with Fura-2-AM (2 µM final concentration) plus Pluronic F-127 (0.1%) for 60 min at room temperature in the dark. Cells were then washed and Fura-2-AM allowed to de-esterify for 60 min in Ca^2+^-free KRG buffer containing 1 mM probenecid. After a final wash, cells were resuspended in KRG buffer with 1.3 mM CaCl₂ at 1×10⁶ cells/mL and plated at 90 µL per well on poly-D-lysine–coated black, clear-bottom 96-well plates.

Fluorescence was recorded at 37 °C on a multimode plate reader using dual-excitation at 340/380 nm and emission at 510 nm. Baseline was acquired for 30 s, after which 10× ligand stocks (10 µL) were added to reach the indicated final concentrations. Where indicated, cells were preincubated for 5 min with the FFA2R antagonist CATPB, Gα_q_ inhibitor YM-254890, or allosteric ligands before agonist addition. Maximal and minimal ratios were determined at the end of each run by sequential addition of ionomycin (0.5 µM final) and EGTA/EDTA to chelate extracellular Ca^2+^. Signals are presented as the 340/380 fluorescence ratio and analyzed as peak over baseline. All measurements were performed in duplicate or triplicate wells per condition and repeated in independent biological replicates.

### BRET data analysis

BRET ratios (BRET²) were calculated as the ratio of light emitted by the acceptor (515 nm) over the donor (410 nm). Ligand-induced ΔBRET^2^ values were obtained by subtracting the BRET ratio of vehicle-treated cells from that of ligand-treated cells. For experiments involving preincubation with allosteric ligands (Cmp58 or AZ1729), ΔBRET^2^ values were calculated by correcting for both baseline vehicle signals and responses to each ligand alone, ensuring quantification of the specific allosteric effect ^31^.

All data were analyzed using GraphPad Prism (version 10.3, GraphPad Software, San Diego, CA, USA). Concentration–response curves were fitted using nonlinear regression (three-parameter fit), and results are presented as mean ± SEM from at least three independent experiments with two technical replicates each. Statistical analyses were performed using one-way ANOVA with Dunnett’s multiple-comparison test or Student’s t-test, as appropriate. Statistical significance was defined as p < 0.05. One star (*) indicates a p-value <0.05, two stars (**) indicates p-value <0.01, and three stars (***) indicates p-value <0.001. If nothing else is stated, the data is considered non-significant (ns). Full analytical procedures and statistical rationale are described in our previous work ^31^.

### Power calculation

#### BRET

An *a priori* power analysis was conducted using G*Power (version 3.1.9.7) to verify that the number of replicates was adequate for detecting a biologically meaningful effect in FFA2R BRET signaling assays. Based on pilot data and prior reports ^31,59^, an expected mean change of 0.10 BRET ratio units with a standard deviation of 0.02 was used. A two-tailed *t*-test (α= 0.05, power = 0.80, equal group sizes) indicated that n= 3 biological replicates per condition provided an actual power of >0.99. To ensure reproducibility, each ligand stimulation was performed in duplicate wells within the same plate. The ebBRET platform used in this study has previously been validated for high sensitivity and low variability, supporting reliable detection of FFA2R-dependent G-protein activation and β-arrestin recruitment with small sample sizes.

#### ROS production

A priori power analyses were performed in G*Power (version 3.1.9.7) for a paired t-test (α = 0.05, power = 0.80). Pilot runs showed low within-donor variability in ROS (small *SD*_difference_), and the antagonist was expected to produce ≥90% inhibition relative to propionate (±PAM). Under these conditions, the standardized paired effect size was estimated at *d_z_* ≥ 2.2 (one-tailed, decrease specified a priori), yielding a required sample size of n= 3 donors. The instrument resolution (1 Mega counts per minute (Mcpm)) is well below the expected effect magnitude, ensuring adequate detectability. For a conservative two-tailed analysis, the corresponding threshold is *d_z_* ≈ 3.0; our pilot variance indicates this criterion is still met. For HL60 independent-group comparisons, the same logic applies using an unpaired t-test, with low pooled SD.

## Declarations

### Ethics approval and consent to participate

This study was conducted at the Sahlgrenska Academy, University of Gothenburg, Sweden. Buffy coats were obtained from the blood bank at Sahlgrenska University Hospital, Gothenburg, Sweden. According to Swedish legislation (Section code 4§ 3p, SFS 2003:460; Lag om etikprövning av forskning som avser människor), no ethical approval was required as the buffy coats were provided anonymously and could not be traced back to specific donors.

### Data and materials availability

All data supporting the conclusions of this study are included in the article and/or its Supplementary Materials. Some of the biosensors used in this study are protected by patents but are available for academic research under a standard material transfer agreement (MTA) upon reasonable request to Michel Bouvier. All other materials are available from the corresponding author upon reasonable request.

### Authors’ contributions

Simon Lind (S.L.) conceived the study, designed and performed all experiments, analyzed the data, prepared the figures, and wrote the manuscript. Ayaan Abdi Ali (A.A.A.) assisted with cell culture, transfections, and data analysis. Sarah Al Hamoud Al Asswad (S.A.H.A.A.) assisted with experimental design and data analysis. Volker Lauschke (V.M.L.) provided critical input on data interpretation and wrote the manuscript. Huamei Forsman (H.F.) contributed to experimental design and provided expertise in neutrophil biology. Claes Dahlgren (C.D.) contributed to study design and assisted in writing the first draft of the manuscript. Linda Johansson (L.C.J.) supervised the project, contributed to study design, interpreted the data and wrote the manuscript. All authors contributed to the writing process, provided feedback on the manuscript, and approved the final submitted version.

## Acknowledgements

We thank Michel Bouvier for providing biosensor constructs and technical expertise in BRET assay methodology.

