## Supplementary material for "Free Fatty Acid Receptor 2 Allosterism is Defined By Cellular Context"

Supplementary figure 1, Lind *et al*

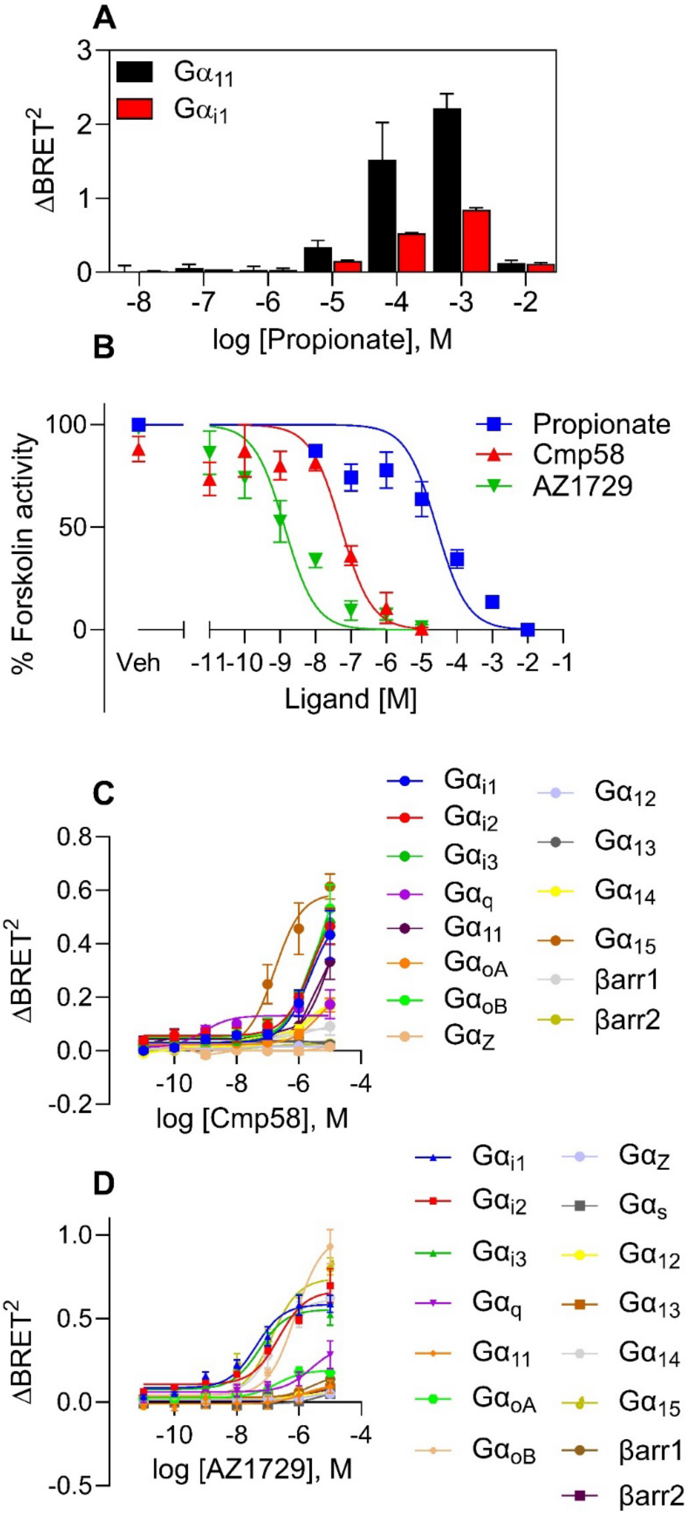


**Supplementary Figure 1**: **Assessment of FFA2R-mediated cAMP inhibition and G protein/β-arrestin activation by allosteric ligands.**

(**A**) Dose dependent activation by propionate until concentration causes acidification effect (-2M) following G$\alpha_{11}$ (black bar) and G$\alpha_{i1}$ activation measured in $\Delta$BRET$^2$ and presented as a bar graph. The bar graph represents data from three independent experiments (mean+SEM); $n$ = 3. (**B**) HEK293T cells transiently expressing FFA2R were stimulated with increasing concentrations of propionate (blue squares), Cmp58 (red triangles), or AZ1729 (green inverted triangles). Intracellular cAMP levels were measured using the GloSensor assay following stimulation with forskolin (10 μM). All ligands produced concentration-dependent inhibition of forskolin-stimulated cAMP accumulation, consistent with FFA2R coupling to G$\alpha_{i/o}$. Data are normalized to forskolin inhibition (in %) and represent mean ± SEM from three independent experiments performed in duplicate. Concentration–response curves of FFA2R allosteric agonists Cmp58 (**C**) and AZ1729 (**D**) monitoring the activation profile of G protein and β-arrestin pathways. The ebBRET methodology was used with different biosensors detecting movement to the plasma membrane following ligand addition, using rGFP-CAAX as the membrane anchor. **(C-D)** subfigures present nonlinear fit dose concentration of the compound in log (molar) on the x-axis and the unitless $\Delta$BRET$^2$ signal on the y-axis in ± SEM with three biological replicates performed in duplicates (n=3).

Supplementary figure 2, Lind *et al*

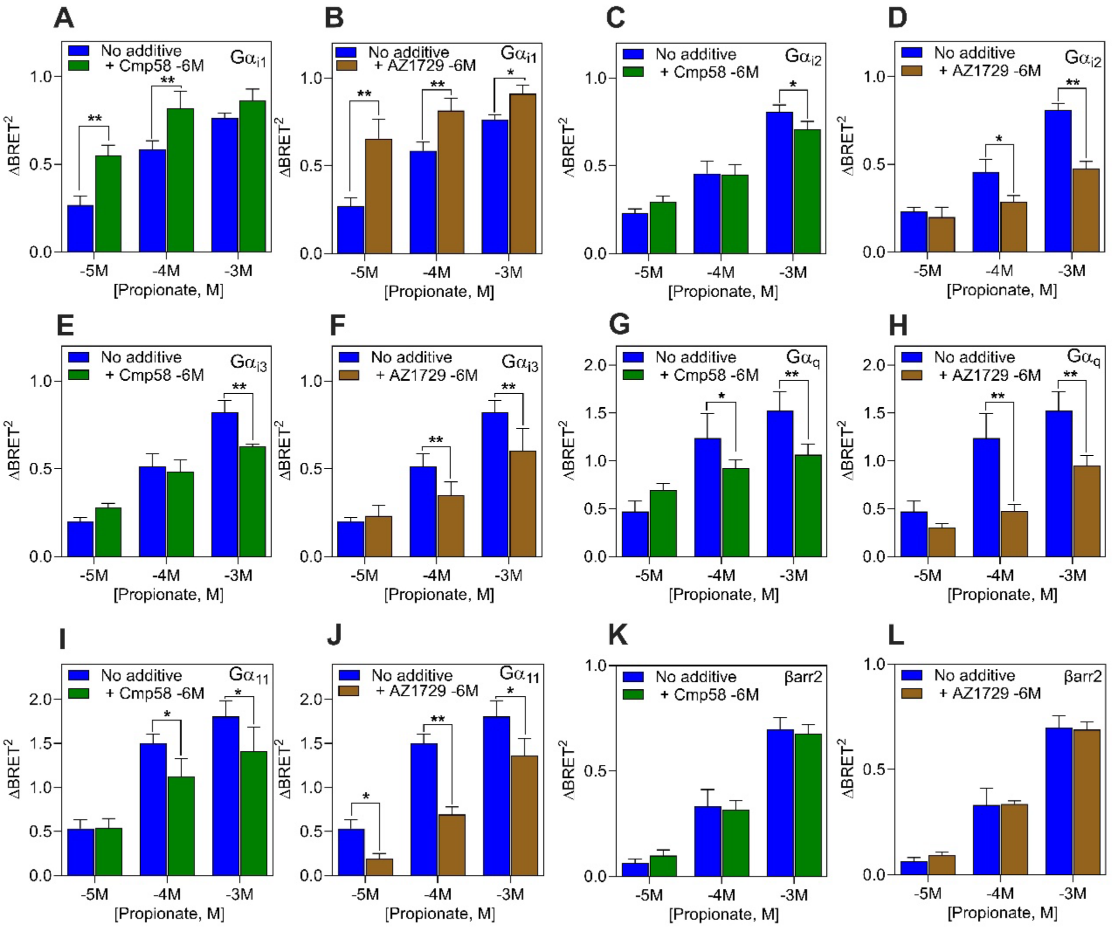


**Supplementary Figure 2: Cmp58 and AZ1729 potentiate propionate induced G$\alpha_{i1}$ activation without affecting β-arrestin 2 recruitment**

For all panels, $BRET^2$ values were first converted to $\Delta BRET^2$ by subtracting the $BRET^2$ of vehicle-treated cells from that of ligand-treated cells. In panels where ligands were preincubated (A–J), a net effect of the preincubated ligand was additionally calculated by subtracting the $\Delta BRET^2$ elicited by the preincubated ligand alone (e.g., 1 µM Cmp58 or AZ172) from the $\Delta BRET^2$ elicited by the combination (*see Materials and Methods for details*). Bar graph with different concentrations of propionate (blue) in the absence or presence of Cmp58 (1 µM, 5-min preincubation; green) or AZ1729 (1 µM, 5-min preincubation; brown). Propionate-induced activation was measured for G$\alpha_{i1}$ (**A–B**), G$\alpha_{i2}$ **(C–D),** G$\alpha_{i3}$ (**E–F**), G$\alpha_q$ (**G–H**), G$\alpha_{11}$ (**I-J**) and recruitment of β-arrestin2 (**K–L**). $\Delta BRET^2$ responses with and without Cmp58/AZ1729 were compared using paired Student's *t*-test. The bar graph represents data from three independent experiments (mean+SEM); $n = 3$.

Supplementary figure 3, Lind *et al*

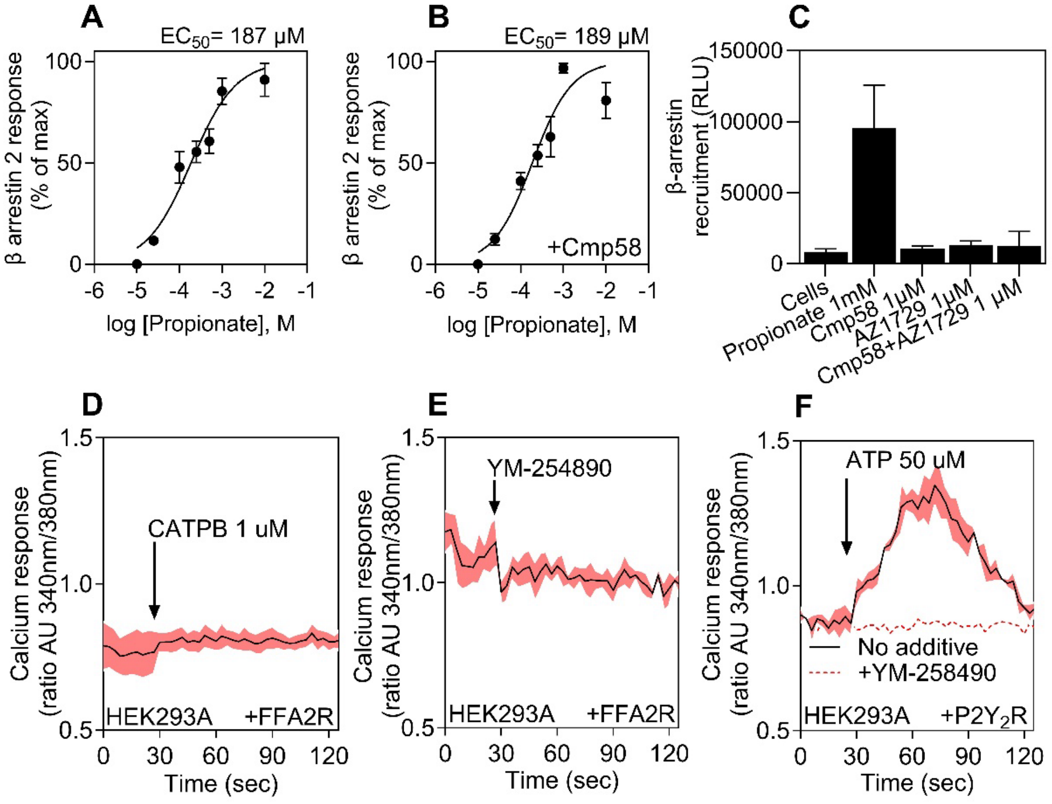


**Supplementary Figure 3**: **Propionate dose dependently recruits β-arrestin 2 but is not affected by the addition of allosteric agonist Cmp58**

(**A**) Concentration-response curves of FFA2R endogenous ligand propionate monitoring the recruitment of β-arrestin2 using NanoBiT assay in HEK293T cells expressing FFA2R. $EC_{50}$-value determined based on the peak relative light units (RLU) responses. Curve fitting was performed by non-linear regression using the sigmoidal dose-response equation (variable-slope). (**B**) As in (A) but monitoring the concentration-response curves of FFA2R endogenous ligand propionate in the presence of a fixed concentration of Cmp58 (1 µM) **C**) The recruitment β-arrestin2 following ligand addition was determined by the production of light (RLU) in HEK293T cells (+FFA2R) alone or cells activated with propionate (1 mM), Cmp58 (1 µM), AZ1729 (1 µM) or Cmp58+AZ1729 (1 µM) using a NanoBiT assay. Peak activities were determined from the peak RLU activities 2 min after ligand addition (mean+SEM, n = 3). **(D–F)** The transient rise of intracellular $Ca^{2+}$ concentration was followed in HEK293T cells transiently expressing FFA2R or $P2Y_2R$ as indicated. (**D**) Cells were stimulated with the FFA2R antagonist CATPB (1 µM). (**E**) Cells were stimulated with the $G\alpha_q$ inhibitor YM-254890 (200 nM). (**F**) Cells transiently expressing $P2Y_2R$ were stimulated with ATP (50 µM) in the absence (solid line) or presence (dashed line) of YM-254890 (200 nM, preincubated for 5 min). Ligand addition is indicated by arrows. Data are representative of three independent experiments, and shaded areas represent the standard error of the mean. Abscissa, time of study (sec); ordinate, increase in intracellular $Ca^{2+}$ $[Ca^{2+}]_i$ given as the change in the ratio between Fura-2 fluorescence at 340 and 380 nm (AU, arbitrary units).
